# Cell signaling coordinates global Polycomb Repressive Complex 2 recruitment and gene expression in murine embryonic stem cells

**DOI:** 10.1101/597674

**Authors:** Mohammad B. Aljazi, Yuen Gao, Yan Wu, George I. Mias, Jin He

## Abstract

The recruitment of Polycomb repressive complex 2 (PRC2) to gene promoters is critical for its function in repressing gene expression in murine embryonic stem cells (mESCs). However, previous studies have demonstrated although the expression of early lineage-specific genes is largely repressed, the genome-wide PRC2 occupancy is unexpectedly reduced in naïve mESCs. In this study, we provide evidence to show the FGF/ERK signaling determines the global PRC2 occupancy through regulating the expression of PRC2-recruting factor JARID2 in naïve mESCs. At the transcriptional level, the de-repression of bivalent genes is predominantly determined by the presence of cell signaling-associated transcription factors but not the status of PRC2 occupancy at gene promoters. Hence, this study not only reveals a key molecular mechanism by which the FGF/ERK signaling in regulating the PRC2 occupancy in mESCs, but also elucidates a fundamental question regarding the functional roles of transcription factors and Polycomb-mediated epigenetic mechanisms in transcriptional regulation.

## Introduction

Polycomb repressive complexes (PRCs) are critical for maintaining mESCs in an undifferentiated state by repressing lineage-specific gene expression (Boyer et al. 2006; Aloia, Di Stefano, and Di Croce 2013; Lee et al. 2006). Previous studies have shown that both PRC1 and PRC2 are recruited to gene promoters and mediate monoubiquitylation of histone H2A lysine 119 (H2AK119ub1) and trimethylation of histone H3 lysine 27 (H3K27me3) respectively (Cao et al. 2002; Wang et al. 2004; Di Croce and Helin 2013), which are essential for their function in repressing gene expression (Blackledge et al. 2019; Tamburri et al. 2019; Cao et al. 2002).

Although the genome-wide binding sites of PRCs in mESCs have been well documented, the molecular mechanisms for recruiting PRCs to specific genomic loci in mammalian cells have not been fully elucidated (Laugesen, Hojfeldt, and Helin 2019). A variety of factors, such as JARID2 and MTF2, have been reported to mediate the PRC2 recruitment (Landeira et al. 2010; Pasini et al. 2010; Shen et al. 2009; Casanova et al. 2011; Li et al. 2017; Oksuz et al. 2018). On the other hand, histone lysine demethylase 2b (KDM2B) recruits a non-canonical PRC1.1 variant to the unmethylated CpG islands (CGIs) in mammalian cells through its CxxC-zinc finger (CxxC-ZF) domain (Wu, Johansen, and Helin 2013; He et al. 2013; Farcas et al. 2012). Recently, KDM2B was also reported to recruit PRC2 though the interaction of JARID2/PRC2 with KDM2B/PRC1.1-mediated histone H2AK119ub1 modification (Cooper et al. 2014).

The repression of differentiation gene expression is essential for mESCs to maintain their pluripotency (Smith 2001). However, differentiation-inducing signals such as the extracellular signal-regulated kinase (ERK) signaling triggered by fibroblast growth factors (FGFs) in serum induce stochastic expression of early lineage-specific genes. After being cultured in 2i serum-free medium in which the FGF/ERK signaling is inhibited and the Wnt/beta-catenin signaling is activated, mESCs turn into a naïve state with reduced basal expression of early lineage-specific genes (Ying et al. 2008; Silva et al. 2008). Along with the gene expression changes, naïve mESCs display two major genome-wide epigenetic changes (Takahashi, Kobayashi, and Hiratani 2018): (i) a global reduction of DNA methylation mainly due to decreased expression of *Uhrf1* and loss of DNA methylation maintenance (Leitch et al. 2013; von Meyenn et al. 2016), and (ii) both genome-wide PRC2 occupancy and its mediated histone H3K27me3 modification are drastically reduced (Marks et al. 2012; Guo et al. 2016; Galonska et al. 2015). Although the underlying molecular mechanisms for the global reduction of PRC2 occupancy are unclear, several recent studies suggest the decreased PRC2 occupancy at CGIs is due to the relocation of PRC2 to the new DNA demethylated non-CGI regions (Walter et al. 2016; van Mierlo et al. 2019; Kumar and Elsasser 2019), or the deficiency of ERK signaling leading to reduced chromatin accessibility to PRC2 (Tee et al. 2014).

To examine the molecular mechanisms leading to the reduced PRC2 occupancy and H3K27me3 modification in naïve mESCs, we performed RNA-sequencing (RNA-seq) analyses to compare the gene expression in wild-type mESCs cultured in serum versus 2i medium. The result showed that the expression of a known PRC2 recruiting factor JARID2 was significantly reduced in naïve mESCs. Reactivation of FGF/ERK signaling increased the expression of *Jarid2*, suggesting the FGF/ERK signaling positively regulated its expression. Similarly, genetic deletion of ERK signaling molecules ERK1 and ERK2 reduced the expression of *Jarid2*, which could be rescued by ectopic expression of wild-type ERK2. ChIP-seq analyses showed that the global occupancy of JARID2/EZH2 and histone H3K27me3 modification were largely reduced at CGIs in both naïve and *Erk1/Erk2* double knockout (*Erk1/Erk2*-dKO) mESCs, which correlated with a global reduction of JARID2 occupancy. Importantly, ectopic expression of *Jarid2* fully restored the global EZH2 occupancy and the H3K27me3 modification at CGIs in both naïve and *Erk1/Erk2*-dKO mESCs. At the transcriptional level, although the PRC2 occupancy and H3K27me3 modification were reduced at bivalent promoters, the FGF/ERK-regulated lineage-specific genes remained silenced while the Wnt signaling-regulated genes were de-repressed in naïve mESCs, suggesting the presence of transcription factors, but the not the status of PRC2 occupancy, played a predominant role in determining transcriptional activation. Thus, this study not only revealed a key molecular mechanism by which the FGF/ERK signaling regulated the PRC2 occupancy in mESCs, but also elucidated a fundamental question regarding the functional roles of transcription factors and Polycomb-mediated epigenetic mechanisms in transcriptional regulation.

## Results

### The *Jarid2* expression is significantly reduced in naïve mESCs

The reduced global PRC2 occupancy in naïve mESCs could be caused by the reduced expression of PRC2 components. To investigate this possibility, we performed RNA-seq analyses to examine the differential gene expression between wild-type E14 mESCs cultured in serum-containing medium (ESC-S) and 2i medium (ESC-2i). The results identified 1905 upregulated genes and 2371 downregulated genes in ESC-2i respectively (cutoff: 1.5 fold expression change and q < 0.05) (Fig. 1A, 1B). Consistent with previous reports, the expression of *Nanog, Klf4*, and *Prdm14* was found to be upregulated in ESC-2i, validating the proper 2i culture condition and the naïve state of mESCs in this study (Fig. 1C) (Guo et al. 2009; Yamaji et al. 2013; Munoz Descalzo et al. 2012). Among all PRC1 and PRC2 core genes, *Jarid2* expression was found to be significantly reduced in mESC-2i compared to that in mESC-S (Fig. 1D, 1E), which was further confirmed by both quantitative real-time PCR (qRT-PCR) and western blot (WB) analyses at the mRNA and protein levels (Fig. 1F, 1G). *Eed*, to a less extend, also showed reduced expression in mESC-2i (Fig. 1D). To corroborate our findings, we re-analyzed the expression of *Jarid2* and *Eed* in naïve mESCs from a published RNA-seq dataset (Marks et al. 2012). Consistent with our findings, *Jarid2* expression was largely reduced in both E14 and TNGA mESC lines cultured in 2i medium (Fig. S1A). However, the change of *Eed* expression varied in these two mESC lines (Fig. S1B). Therefore, the observed reduced *Eed* expression in the E14 mESC line was likely to be a cell line-specific effect. The expression of other PRC1 and PRC2 core genes including *Kdm2b, Ring1b, Ezh1, Ezh2, Suz12*, and *Mtf2* did not show statistically significant difference between mESC-S and mESC-2i (Fig.S1C, D). Based on these results, we concluded that the expression of PRC2 recruiting factor JARID2 was significantly reduced in naïve mESCs.

**Figure 1.**
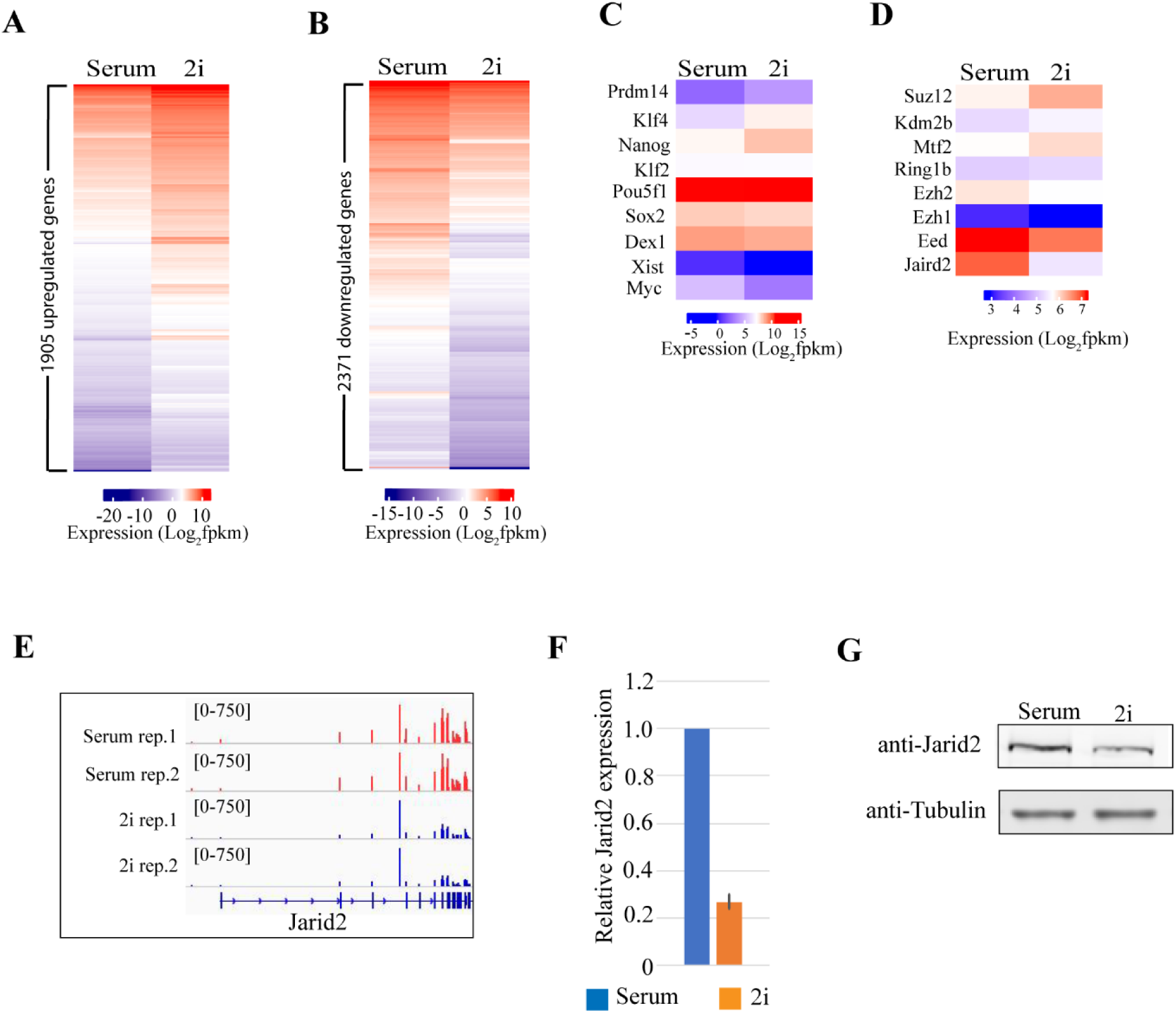
The *Jarid2* expression is significantly reduced in naïve mESCs. (**A**) Heatmap showing 1905 genes upregulated in mESC-2i. (**B**) Heatmap showing 2371 genes downregulated in mESC-2i. (**C**) Heatmap showing the expression of pluripotent genes in mESC-S and mESC-2i. (**D**) Heatmap showing the expression of Polycomb core genes in mESC-S and mESC-2i. The expression of Jarid2 was significantly reduced in mESC-2i. (**E**) The IGV genome browser view of *Jarid2* expression in mESC-S and mESC-2i. (**F**) qRT-PCR analysis showing *Jarid2* had a reduced mRNA level in mESC-2i. The results were normalized against levels of *Gapdh* and the expression level in mESC-S was arbitrarily set to 1. The error bar represents the standard deviation (n=3). (**G**) Western blot analysis showing JARID2 had a reduced protein level in mESC-2i.

### FGF/ERK signaling positively regulates the *Jarid2* expression

The FGF/ERK signaling is blocked by removal of serum and supplement of MEK inhibitor PD0325901 in 2i medium. To examine whether the reduced expression of *Jarid2* is caused by the deficiency of FGF/ERK signaling in 2i medium, we added the fetal bovine serum back to, and removed the MEK inhibitor PD0325901 from the 2i medium. RNA-seq analyses were performed to examine the gene expression at different time points after the medium change (Fig. 2A). The activation of FGF/ERK signaling was monitored by WB analyses on the phosphorylated MAPK p42/44. The result showed that the phosphorylated MAPK p42/44 were not detected in ESC-2i. However, the phosphorylated proteins quickly increased and reached to a constant level at 6 hours after the medium switch, confirming the reactivation of FGF/ERK signaling (Fig. 2B). The RNA-seq analyses at 0, 6 and 12 hours after the medium change identified 309 genes upregulated in response to the activation of FGF/ERK signaling (Fig. S2A). Kyoto Encyclopedia of Genes and Genomes (KEGG) pathway analysis revealed that the upregulated genes were enriched in MAPK signaling pathway, signaling involved in regulating stem cell pluripotency, signaling pathway in cancer, and HIF signaling pathway (cutoff: Benjamini adjusted p < 0.05), further confirming that the FGF/ERK signaling pathway was properly activated in mESCs after the medium change (Fig. S2B, Table S1). Importantly, RNA-seq results showed that *Jarid2* expression is increased in response to the reactivation of FGF/ERK signaling (Fig.2C). The increased *Jarid2* expression was further confirmed by qRT-PCR and WB analyses at both mRNA and protein levels (Fig. 2D, 2E). Based on these results, we concluded that the FGF/ERK signaling positively regulated the expression of *Jarid2* and blocking of this signaling pathway in 2i medium led to its reduced expression at both mRNA and protein levels in naïve mESCs.

**Figure 2.**
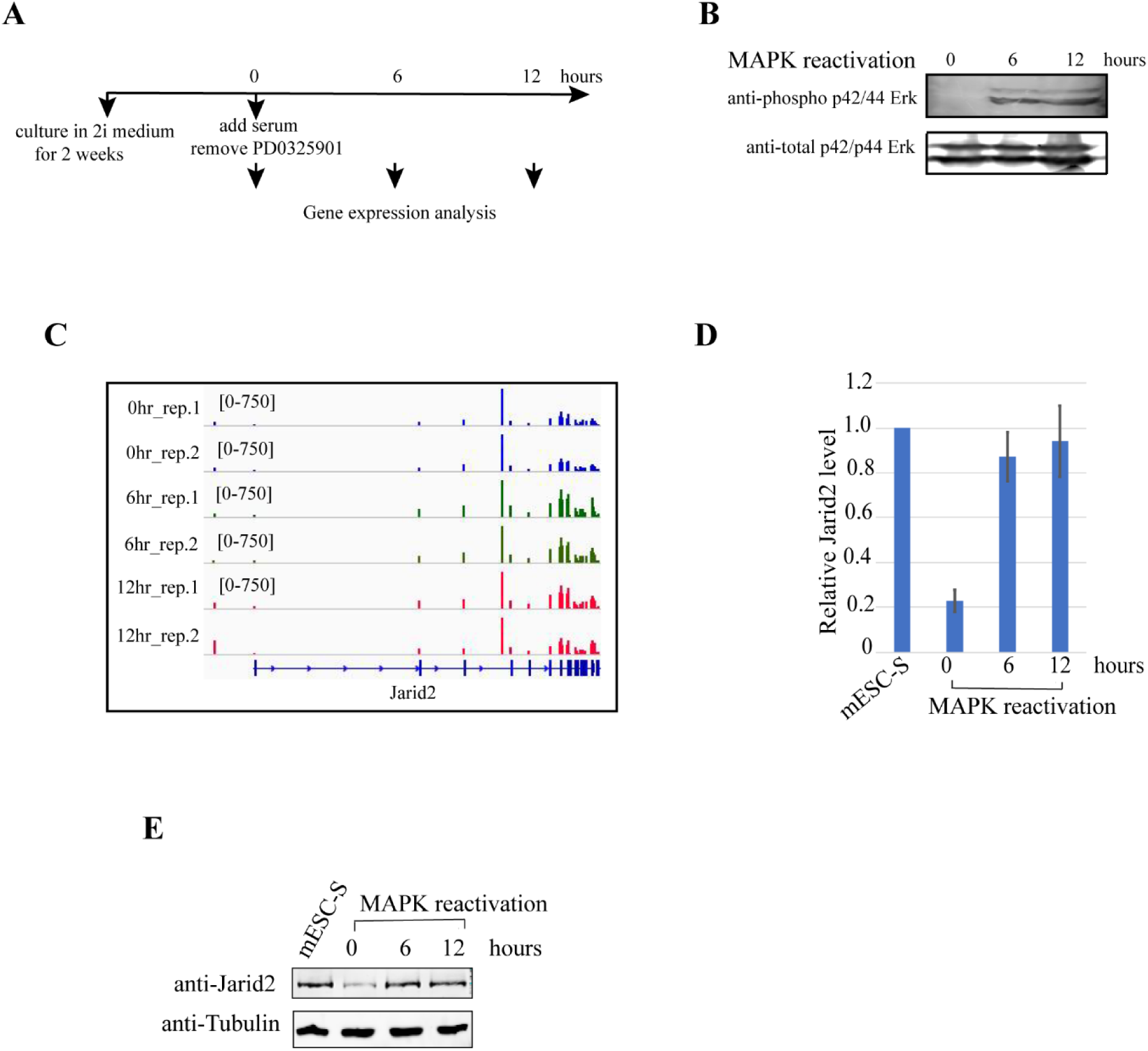
FGF/ERK signaling positively regulates the *Jarid2* expression. (**A**) Schematic chart showing the experimental design for the analysis of gene expression in response to the reactivation of FGF/ERK signaling. (**B**) Western blot analysis showing phosphorylated p42/44 MAPKs were increased in response to the reactivation of FGF/ERK signaling. (**C**) The IGV genome browser view of *Jarid2* expression before, 6 hours and 12 hours after the reactivation of FGF/ERK signaling. (**D**) qRT-PCR analysis showing the *Jarid2* expression was upregulated at the mRNA level in response to the reactivation of FGF/ERK signaling. The results were normalized against levels of *Gapdh* and the expression level in mESC-S is arbitrarily set to 1. The error bars represent the standard deviation (n=3). (**E**) Western blot analysis showing JARID2 expression was upregulated at the protein level in response to the reactivation of FGF/ERK signaling in mESC-S and different time points after reactivation of FGF/ERK signaling.

### Knockout of *Erk1/Erk2* reduces the expression of *Jarid2* in mESCs cultured in serum-containing medium

A previous study reported the genetic depletion of MAPK signaling molecules ERK1 and ERK2 in mESCs led to a reduction of PRC2 occupancy and H3K27me3 modifications at bivalent promoters (Tee et al. 2014). Since the study did not examine the *Jarid2* expression in the *Erk1/Erk2*-depleted mESCs, we asked whether the observed reduction of JARID2 occupancy at bivalent promoters in the *Erk1/Erk2*-depleted mESCs is due to reduced *Jarid2* expression after the disruption of ERK signaling as observed in naïve mESCs.

To examine this possibility, we established multiple *Erk1/Erk2*-dKO mESC lines by the Crispr-Cas9-mediated gene knockout approach. The depletion of ERK1/ERK2 proteins in two independent *Erk1/Erk2*-dKO lines were confirmed by WB analyses (Fig. 3A, lane 1-3). Compared to wild-type mESCs, the *Erk1/Erk2*-dKO mESCs cultured in the serum-containing medium supplemented with a GSK3 inhibitor CHIR99021 maintain relatively normal colony morphology (Fig. S3A). RNA-seq analysis showed that similar to mESC-2i, the *Erk1/Erk2*-dKO mESCs had increased expression of *Prdm14, Nanog*, and *Klf4*, reduced *c-myc* expression, and similar expression of *Oct4* and *Sox2*, suggesting the *Erk1/Erk2*-dKO mESCs maintained the expression of core pluripotent genes under the experimental condition (Fig. S3B). Importantly, RNA-seq analyses showed that *Jarid2* expression was significantly reduced while other PRC1 and PRC2 core genes had similar or increased expression levels in the *Erk1/Erk2*-dKO mESCs (Fig. 3B, 3C). The decreased *Jarid2* expression in the *Erk1/Erk2*-dKO cells is confirmed by qRT-PCR and WB analyses at the mRNA and protein levels (Fig. 1. lane 1-3, Fig. 3D). To corroborate the findings, we recused the ERK signaling by lentiviral vectors expressing wild-type *Erk2* fused with a puromycin resistant cassette linked by a self-cleaved P2A peptide. After transduction, the transduced cells were selected by puromycin and the expression of exogenous ERK2 is confirmed by WB analysis (Fig. 3A lane 4). RNA-seq analysis showed the *Jarid2* expression is restored in the wild-type *Erk2-*rescued cells (Fig. 3B, 3C), which is further confirmed by qRT-PCR and WB analyses (Fig. 3A lane 4, 3D).

**Figure 3.**
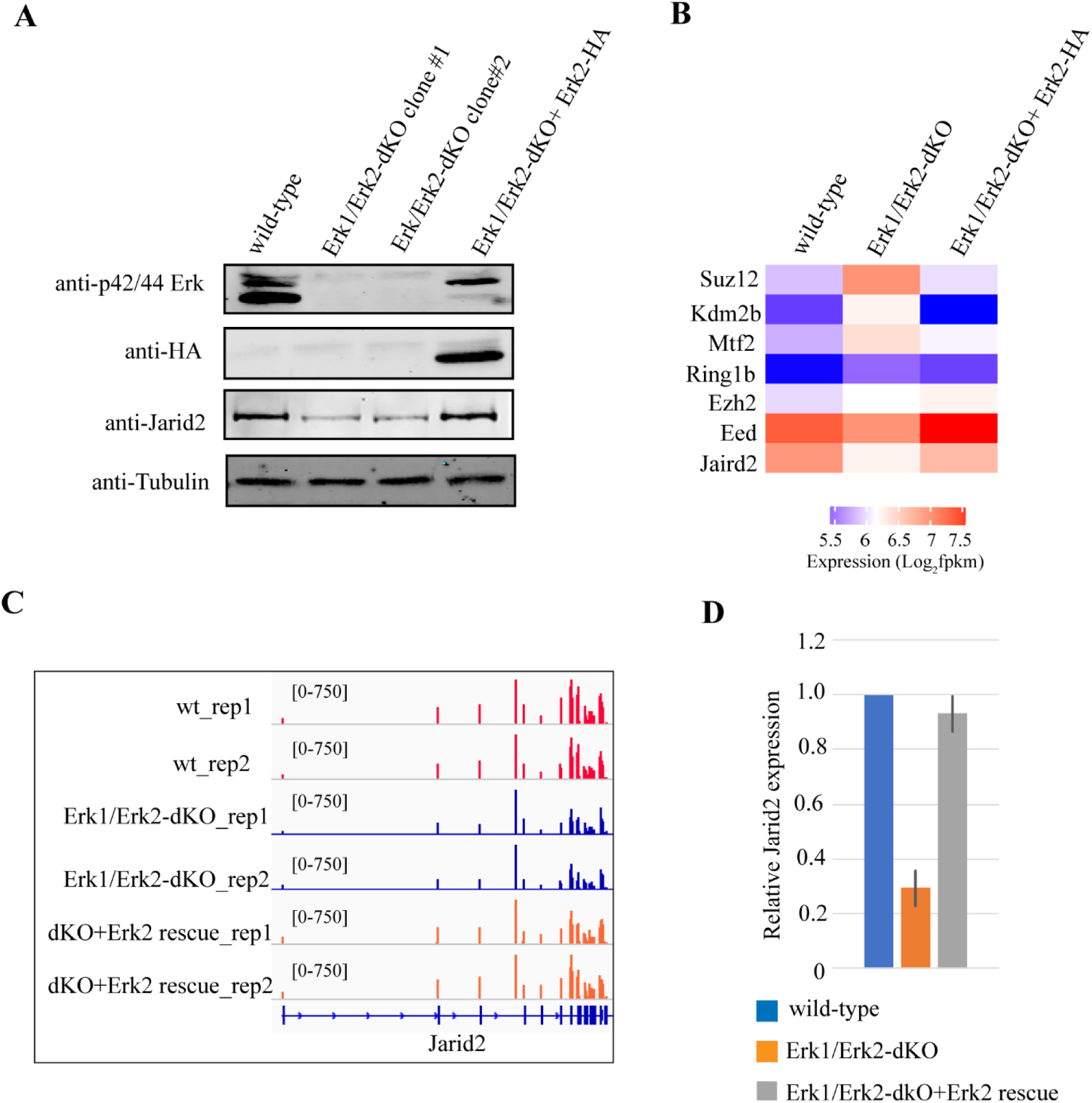
Knockout of *Erk1/Erk2* reduces the *Jarid2* expression in mESCs cultured in serum-containing medium. (**A**) Western blot analysis showing p42/44 MAPKs, ectopic expressed wild-type ERK2-HA, and JARID2 proteins in wild-type, *Erk1/Erk2*-dKO clones, and *Erk1/Erk2*-dKO rescued with wild-type ERK2-HA mESCs. (**B**) Heatmap showing the expression of Polycomb core genes in wild-type, *Erk1/Erk2*-dKO, and *Erk1/Erk2*-dKO rescued with wild-type ERK2-HA mESCs. (**C**) The IGV genome browser view of *Jarid2* expression in in wild-type, *Erk1/Erk2*-dKO, and *Erk1/Erk2*-dKO rescued with wild-type Erk2-HA mESCs. (**D**) qRT-PCR analysis showing the *Jarid2* expression in wild-type, *Erk1/Erk2*-dKO, and *Erk1/Erk2*-dKO rescued with wild-type ERK2-HA mESCs. The results were normalized against levels of *Gapdh* and the expression level in wild-type mESCs was arbitrarily set to 1. The error bars represent standard deviation (n=3).

Based on these results, we concluded that *Jarid2* expression in mESCs was positively regulated by the ERK signaling. Same as the chemical inhibition of FGF/ERK signaling in 2i medium, the genetic disruption of ERK signaling led to reduced *Jarid2* expression in the mESCs cultured in serum-containing medium.

### The genome-wide PRC2 occupancy at CGIs is largely reduced in naïve mESCs

To examine the global epigenetic changes in naïve mESCs, we performed ChIP-seq analyses to compare the genome-wide occupancy of PRC2 components JARID2/EZH2, PRC1 components KDM2B/RING1B, and PRC2-mediated histone H3K27me3 modification as well as Trithorax MLL1/MLL2 complexes-mediated H3K4me3 modification in mESC-S and mESC-2i. Since Polycomb complexes and Trithorax MLL1/MLL2 complexes were globally recruited to CGIs in mammalian cells, we measured the overall Polycomb occupancy and histone H3K27me3/H3K4me3 modifications by calculating the total normalized mapped reads within CGIs and a 10kb-window of CGI-flanking regions. Consistent with previous reports (Marks et al. 2012; Guo et al. 2016), both genome-wide PRC2 occupancy and histone H3K27me3 modification were largely reduced at CGIs in naïve mESCs, which correlated well with the global decreased JARID2 occupancy (Fig. 4A-C, 4E). However, KDM2B, one PRC1.1 component that binds to unmethylated CGIs through its CxxC-ZF domain, had similar genome-wide occupancy at CGIs in mESC-S and mESC-2i (Fig. 4D, 4E). In contrast to the reduced H3K27me3 modification, the global H3K4me3 modification at CGIs was slightly increased in mESC-2i (Fig. S4A). Also, in contrast to the EZH2 occupancy that was reduced at both GGIs and 10-kb CGI-flanking sites, the occupancy of PRC1 core component RING1B was mainly reduced at the CGI flanking regions but not the CGI center (Fig. S4B).

**Figure 4.**
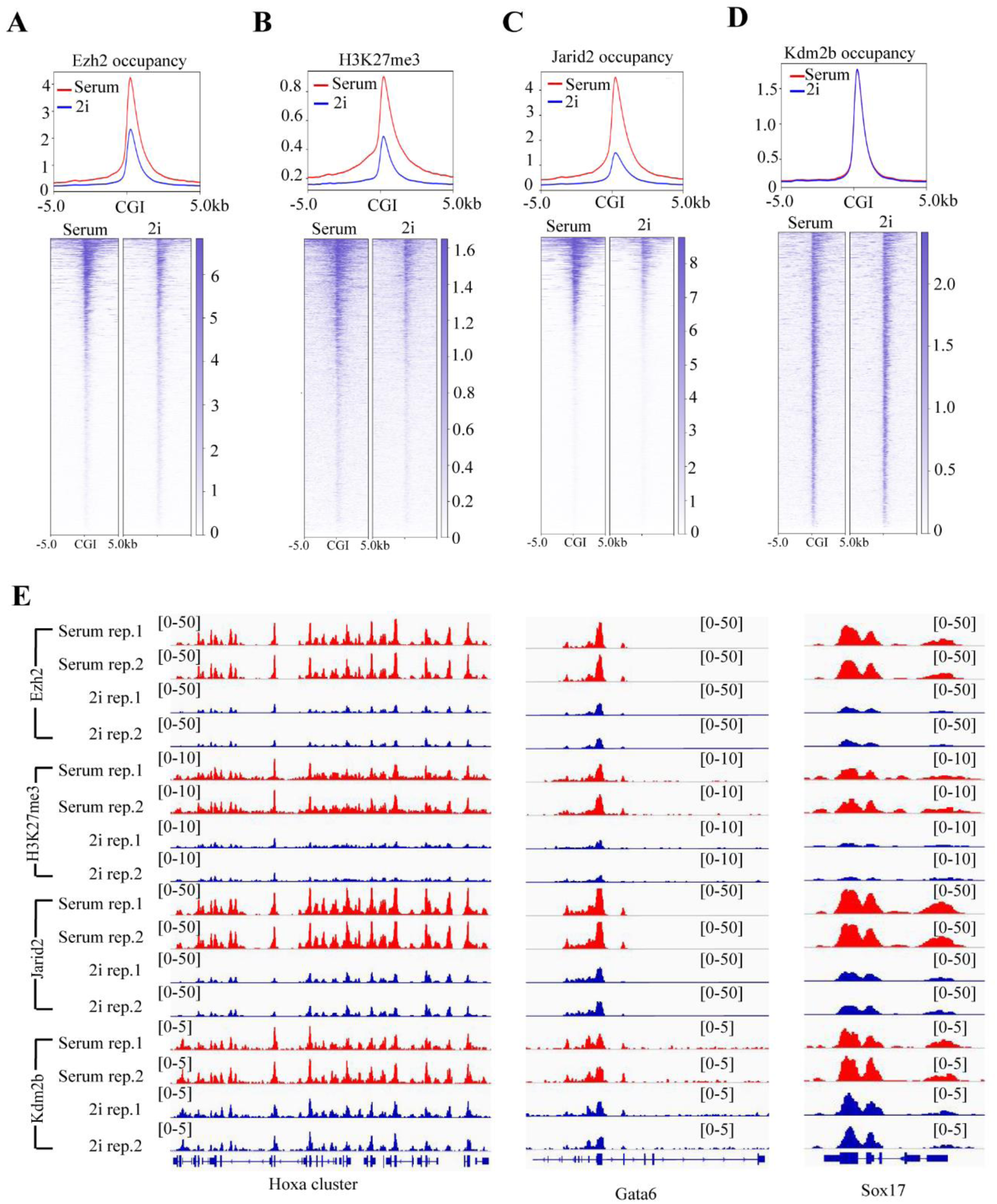
The genome-wide PRC2 occupancy at CGIs is largely reduced in naïve mESCs. (**A**) The plot (upper) and heatmap (bottom) showing the EZH2 occupancy at CGIs and 10kb-window flanking CGIs in mESC-S and mESC-2i. (**B**) The plot (upper) and heatmap (bottom) showing the histone H3K27me3 modification at CGIs and 10kb-window flanking CGIs in mESC-S and mESC-2i. (**C**) The plot (upper) and heatmap (bottom) showing the JARID2 occupancy at CGIs and 10kb-window flanking CGIs in mESC-S and mESC-2i. (**D**) The plot (upper) and heatmap (bottom) showing the KDM2B occupancy at CGIs and 10kb-window flanking CGIs in mESC-S and mESC-2i. (**E**) The IGV genome browser view of EZH2, JARID2, KDM2B occupancy and histone H3K27me3 modification at the representative *Hoxa* gene locus (left panel), *Gata6* (middle panel), and *Sox17* (right panel). The red color represents the level in mESC-S (serum) and the blue color represents the level in mESC-2i (2i).

To examine whether PRC2 occupancy and H3K27me3 modification were relocated to non-CGI regions, we calculated the overall occupancy of JARID2/EZH2 and H3K27me3 modification over extended 50kb CGI-flanking regions. The results did not reveal significant increased PRC2 occupancy and H3K27me3 modification within 50kb CGI-flanking regions (Fig. S4C-E). Since JARID2 is a known PRC2 recruiting factor (Shen et al. 2009; Pasini et al. 2010; Peng et al. 2009), these results suggest that reduced *Jarid2* expression is like to be a key molecular mechanism leading to the reduced global PRC2 occupancy and histone H3K27me3 modification at CGIs in naïve mESCs.

### Ectopic expression of *Jarid2* restores the global PRC2 occupancy at CGIs and bivalent promoters in naïve mESCs

To further examine whether the reduced *Jarid2* expression was the main reason causing the global reduction of PRC2 occupancy at CGIs in naïve mESCs, we rescued the *Jarid2* expression in naïve mESCs by lentiviral viruses expressing *Jarid2* fused with a puromycin resistant cassette linked by a self-cleaved P2A peptide. After transduction, the transduced cells were selected by puromycin in the medium. RNA-seq analyses showed that the *Jarid2* expression increased to a comparable level as that in mESC-S (Fig. S5A). The rescued *Jarid2* expression is confirmed by qRT-PCR and WB analyses at both mRNA and protein levels (Fig. S5B, S5C). The ChIP-seq analyses demonstrated that the global EZH2 occupancy and histone H3K27me3 modification were fully restored, which correlated with the rescued global JARID2 occupancy in naïve mESCs (Fig. 5A-5D). Next, we extracted a total of 2830 unique bivalent promoters with both H3K27me3 and H3K4me3 modifications and analyzed the JARID2/EZH2 occupancy as well as H3K27me3 modifications on these promoters (Fig. S5D). Same as CGIs, the bivalent promoters had reduced JARID2/EZH2 occupancy and H3K27me3 modifications in naïve mESCs, which could be fully rescued by ectopic expression of *Jarid2* (Fig. S5E-S5G). Based on these results, we concluded that the reduced *Jarid2* expression was the main molecular mechanism leading to the reduction of global PRC2 occupancy and histone H3K27me3 modification at CGIs and bivalent domains in naive mESCs.

**Figure 5.**
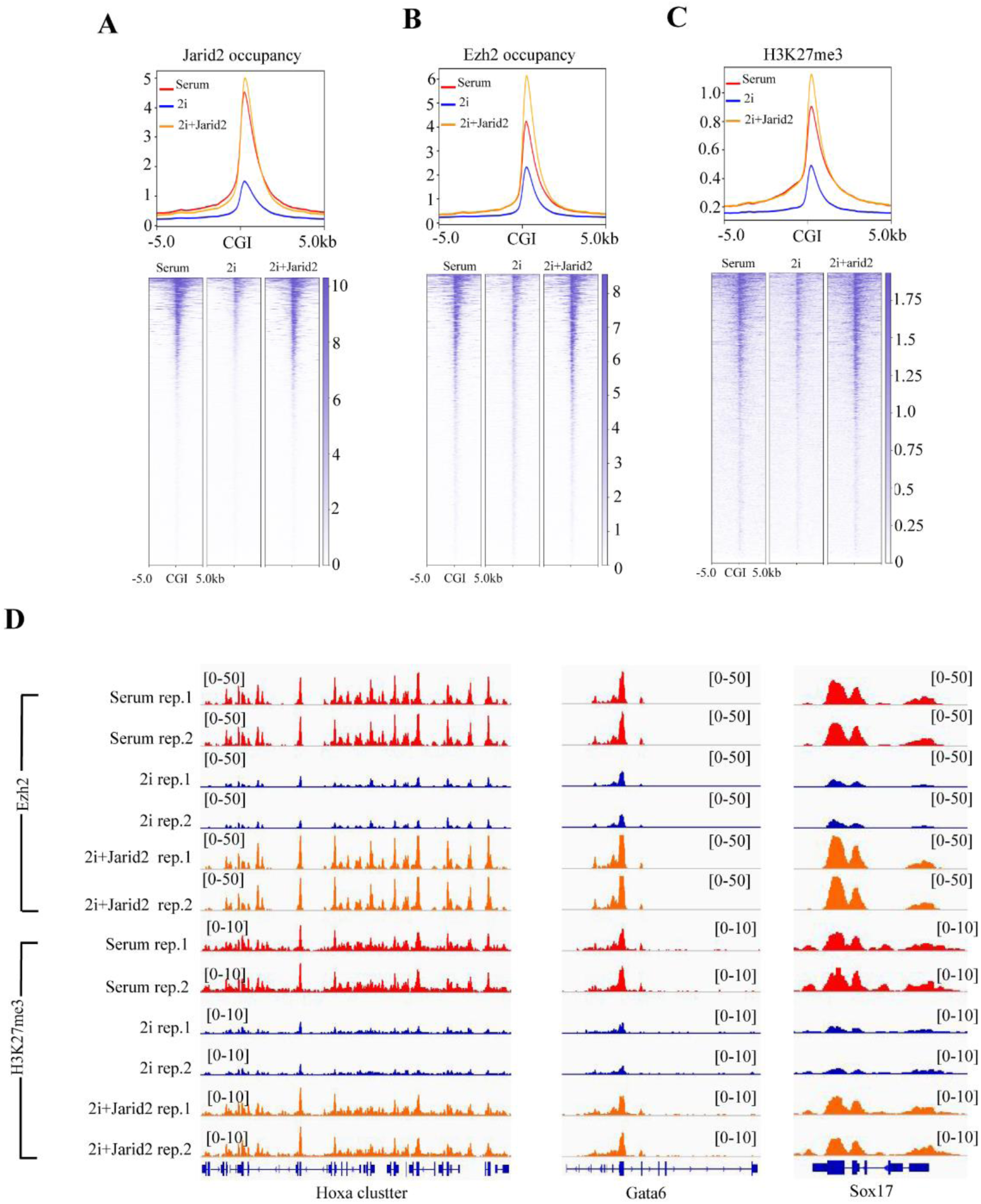
Ectopic expression of *Jarid2* restores the global PRC2 occupancy and histone H3K27me3 modification in naïve mESCs. (**A**) The plot (upper) and heatmap (bottom) showing the JARID2 occupancy at CGIs and 10kb-window flanking CGIs in mESC-S (serum), mESC-2i (2i), and mESC-2i with ectopically expressed *Jarid2* (2i+Jarid2). (**B**) The plot (upper) and heatmap (bottom) showing the EZH2 occupancy at CGIs and 10kb-window flanking CGIs in mESC-S (serum), mESC-2i (2i), and mESC-2i with ectopically expressed *Jarid2* (2i+Jarid2). (**C**) The plot (upper) and heatmap (bottom) showing the histone H3k27me3 at CGIs and 10kb-window flanking CGIs in mESC-S (serum), mESC-2i (2i), and mESC-2i with ectopically expressed *Jarid2* (2i+Jarid2). The red line represents the level in mESC-S (serum), the blue line represents the level in mESC-2i (2i), and the orange line represents the level in mESC-2i with ectopically expressed *Jarid2* (2i+Jarid2). (**D**) The IGV genome browser view of EZH2 and histone H3K27me3 modification at the representative *Hoxa* gene locus (left panel), *Gata6* (middle panel), and *Sox17* (right panel). The red color represents the level in mESC-S (serum), the blue color represents the level in mESC-2i (2i), and the orange color represents the level in mESC-2i with ectopically expressed *Jarid2* (2i+Jarid2).

### Ectopic expression of *Jarid2* restores the global PRC2 occupancy and histone H3K27me3 modification in the *Erk1/Erk2*-dKO mESCs

A previous study reported that genetic depletion of *Erk1/Erk2* resulted in reduced PRC2 occupancy and histone H3K27me3 modification at bivalent promoters in mESCs (Tee et al. 2014). Using the established *Erk1/Erk2*-dKO mESCs (Fig. 3), we performed ChIP-seq analyses to examine the genome-wide occupancy of JARID2/EZH2 and H3K27me3 modifications in these cells. The results showed that the occupancy of JARID2/EZH2 and H3K27me3 modification were largely reduced not only at the bivalent promoters (Fig. S6A-C), but more generally at CGIs (Fig. 6A-C). Importantly, the ectopic expression of wild-type *Erk2* in the *Erk1/Erk2*-dKO mESCs not only largely restored the *Jarid2* expression (Fig. 3), but also rescued the JARID2/EZH2 occupancy and histone H3K27me3 modification at both bivalent promoters and global CGIs (Fig.6A-C, S6A-C).

**Figure 6.**
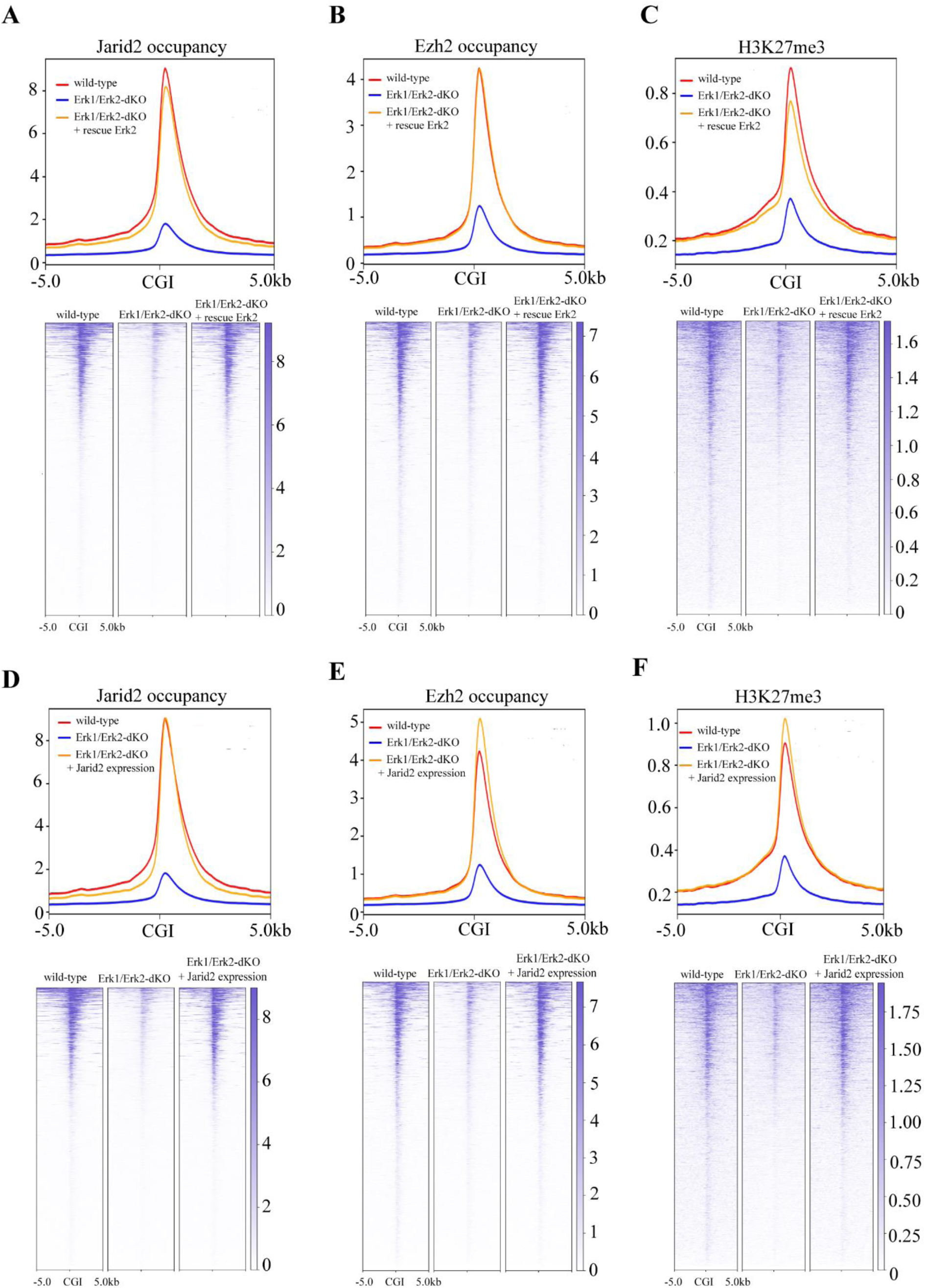
Ectopic expression of *Erk2* or *Jarid2* restores the global PRC2 occupancy and histone H3K27me3 modification at CGIs in *Erk1/Erk2*-dKO mESCs. (**A**) The plot (upper) and heatmap (bottom) showing the JARID2 occupancy at CGIs and 10kb-window flanking CGIs in wild-type mESCs (wild-type), *Erk1/Erk2*-dKO mESCs (*Erk1/Erk2*-dKO), and *Erk1/Erk2*-dKO cells rescued with wild-type *Erk2* (Erk1/Erk2-dKO + rescue *Erk2*). (**B**) The plot (upper) and heatmap (bottom) showing the EZH2 occupancy at CGIs and 10kb-window flanking CGIs in wild-type, *Erk1/Erk2*-dKO, and *Erk1/Erk2*-dKO + rescue *Erk2* mESCs. (**C**) The plot (upper) and heatmap (bottom) showing the histone H3k27me3 modification at CGIs and 10kb-window flanking CGIs in wild-type, *Erk1/Erk2*-dKO, and *Erk1/Erk2*-dKO + rescue *Erk2* mESCs. (**D**) The plot (upper) and heatmap (bottom) showing the JARID2 occupancy at CGIs and 10kb-window flanking CGIs in wild-type mESCs (wild-type), *Erk1/Erk2*-dKO mESCs (*Erk1/Erk2*-dKO), and *Erk1/Erk2*-dKO cells with ectopically expressed *Jarid2* (*Erk1/Erk2*-dKO + J*arid2* expression). (**E**) The plot (upper) and heatmap (bottom) showing the EZH2 occupancy at CGIs and 10kb-window flanking CGIs in wild-type, *Erk1/Erk2*-dKO, and *Erk1/Erk2*-dKO + *Jarid2* expression mESCs. (**F**) The plot (upper) and heatmap (bottom) showing the histone H3k27me3 modification at CGIs and 10kb-window flanking CGIs in wild-type, *Erk1/Erk2*-dKO, and *Erk1/Erk2*-dKO + *Jarid2* expression mESCs.

To further examine whether the reduced JARID2-mediated PRC2 recruitment caused the reduction of JARID2/EZH2 occupancy and H3K27me3 modification at CGIs in the *Erk1/Erk2*-dKO cells, we ectopically expressed *Jarid2* in the *Erk1/Erk2*-dKO cells (Fig. S6D). ChIP-seq analyses showed that the occupancy of JARID2/EZH2 and histone H3K27me3 modification at CGIs as well as bivalent promoters were fully restored in the ectopically *Jarid2*-expressed *Erk1/Erk2*-dKO mESCs (Fig. 6D-E, S6E-G). Thus, same as in the naïve mESCs, the reduced genome-wide PRC2 occupancy at CGIs and bivalent promoters appeared to be mainly caused by the reduced *Jarid2* expression and JARID2-mediated PRC2 recruitment in the *Erk1/Erk2*-dKO mESCs.

### De-repression of bivalent genes appears to be determined by the presence of signaling-associated transcription factors but not the status of PRC2 occupancy in naïve mESCs

To examine whether the reduced PRC2 occupancy led to transcriptional changes of its regulated genes in naïve mESCs, we compared the expression of 2830 bivalent genes in mESC-S versus mESC-2i by RNA-seq analyses. The results identified 589 downregulated genes and 253 de-repressed genes in mESC-2i compared to mESC-S (cutoff: 1.5 fold expression change and q < 0.05) (Fig. 7A, S7A, S7B). Since all bivalent gene promoters had the same reduced PRC2 occupancy and histone H3K27me3 modification in mESC-2i (Fig. S5E-G), the distinct transcriptional status of these two gene groups suggested that the transcriptional activation was determined by factors other than the PRC2 occupancy and its-mediated histone H3K27me3 modification at promoters. The gene ontology (GO) analysis showed that the downregulated genes had enriched GO terms involved in cell differentiation, organ morphogenesis, and multicellular development (cutoff: Benjamini adjusted p < 0.05), which included multiple primitive endoderm-specific genes such as *Gata6, Dab1, Dab2, Sox7*, and *Sox17* that were known to be activated by the FGF/ERK signaling and its associated transcription factors (Fig. 7B, S7C, Table S2). On the other hand, the enriched GO terms of de-repressed genes were involved in multicellular development, transcriptional regulation, and canonical Wnt signaling (cutoff: Benjamini adjusted p < 0.05) (Fig. S7D, Table S3). Notably, multiple known Wnt/beta-catenin direct target genes such as *Axin2, T, Tcf7, Snai1*, and *Cdx1* were de-repressed in naïve mESCs (Roose et al. 1999; Yan et al. 2001; Lustig et al. 2002; ten Berge et al. 2008; Horvay et al. 2011; Lickert et al. 2000), suggesting the de-repression of these genes might be induced by transcription factors associated with the GSK3 inhibitor CHIR99021-activated Wnt/beta-catenin signaling in the 2i medium.

**Figure 7.**
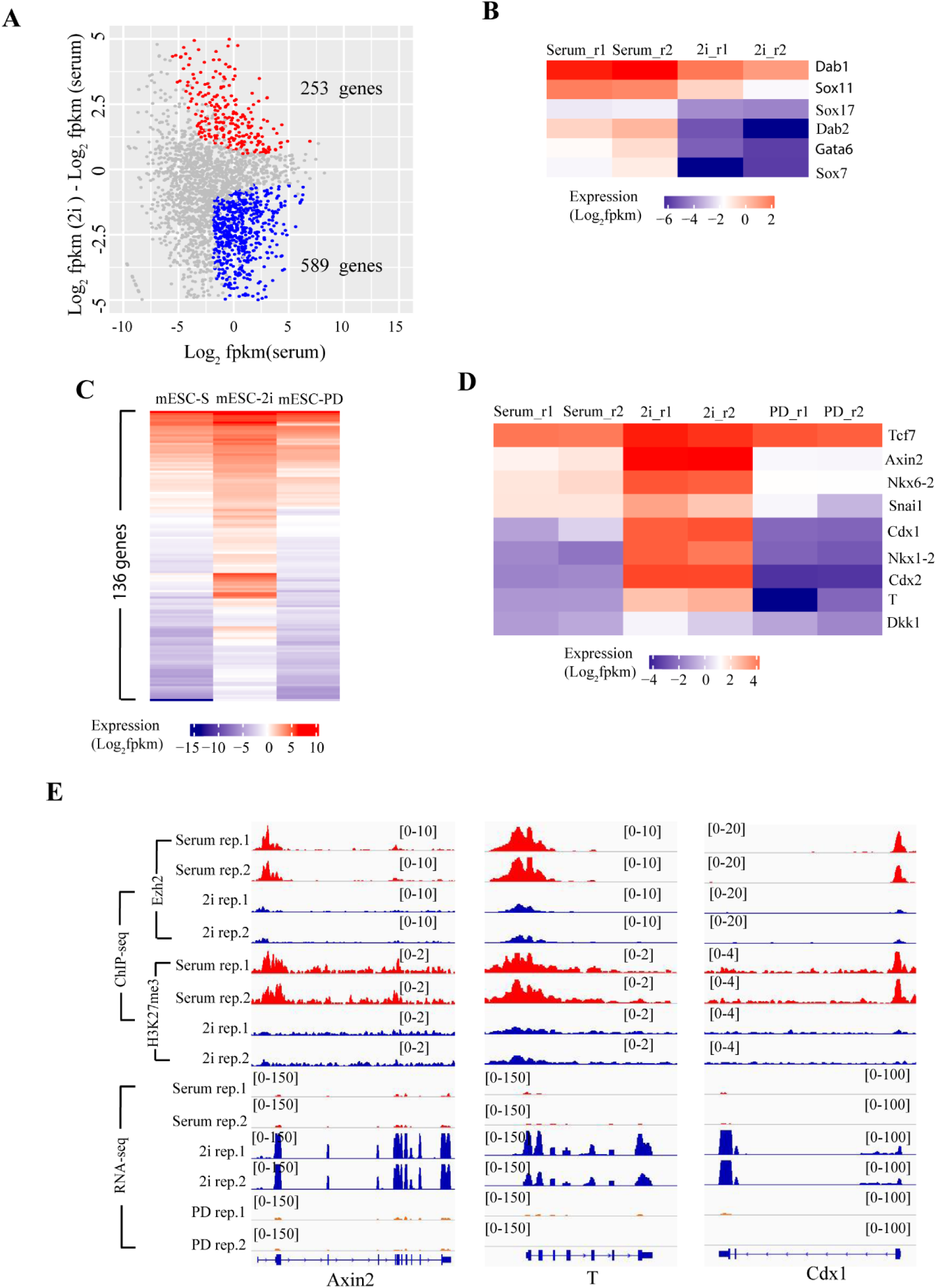
De-repression of bivalent genes is determined by the presence of signaling-associated transcription factors but not the status of PRC2 occupancy in naive mESCs. (**A**) RNA-seq results showing significant 253 up- and 589 down-regulated bivalent genes in mESC-2i compared to that in mESC-S. (**B**) Heatmap showing the expression of representative primitive endoderm-specific genes in mESC-S and mESC-2i. (**C**) Heatmap showing 136 bivalent genes de-repressed in mESC-2i and re-silenced after inactivation of Wnt/beta catenin signaling (mESC-PD). (**D**) Heatmap showing the representative Wnt/beta-catenin direct target genes de-repressed in mESC-2i and re-silenced after inactivation of Wnt/beta catenin signaling (mESC-PD). (**E**) The IGV genome browser view of EZH2 occupancy, H3K27me3 modification, and gene expression of representative Wnt/beta-catenin direct target genes *Axin2, T*, and *Cdx1* in mESC-S, mESC-2i and mESC-PD.

To further examine whether these genes were activated by the Wnt/beta-catenin signaling in naïve mESCs, we inactivated the Wnt/beta-catenin signaling by removing CHIR99021 and leaving a single MEK inhibitor PD0325901 in the medium (mESC-PD). RNA-seq analysis showed that 136 out of 253 genes that were de-repressed in mESC-2i, including all known Wnt/beta-catenin direct target genes such as *Axin2, T, Tcf7, Snai1*, and *Cdx1*, lost the transcriptional de-repression and were re-silenced in mESC-PD (Fig. 7C, 7D), confirming that the de-repression of these genes in mESC-2i depended on the activated Wnt/beta-catenin signaling and its associated transcription factors.

Hence, all these results suggested that the reduced PRC2 occupancy and its mediated histone H3K27me3 modification at gene promoters alone was insufficient to activate transcription of bivalent genes in naïve mESCs. Compared to the reduced PRC2 occupancy at gene promoters, the presence of transcription factors appeared to be necessary and play a predominant role in inducing transcriptional activation.

## Discussion

In mammalian cells PRC1 and PRC2 are recruited to unmethylated CGIs and CGI-associated bivalent gene promoters. The occupancy of Polycomb complexes and their mediated covalent histone modifications at gene promoters are critical for maintaining mESCs in an undifferentiated state by preventing stochastic transcription of lineage differentiation genes.

Previous studies have revealed that both JARID2 and MTF2 are required for the recruitment of PRC2 (Landeira et al. 2010; Pasini et al. 2010; Shen et al. 2009; Casanova et al. 2011; Li et al. 2017; Oksuz et al. 2018), while KDM2B recruits the PRC1.1 variant to unmethylated CGIs through its CxxC-ZF domain that preferentially binds to unmethylated CG-rich DNA sequences (Wu, Johansen, and Helin 2013; He et al. 2013; Farcas et al. 2012). Although the exact molecular mechanisms underlying the PRC2 recruitment to its targets are not fully elucidated, it is believed that the preferential binding to unmethylated CG-rich DNA sequences by the Polycomb recruiting factors JARID2, MTF2, and KDM2B is important to determine the genome-wide Polycomb occupancy at CGIs in mammalian cells (Laugesen, Hojfeldt, and Helin 2019; Deaton and Bird 2011).

Compared to the mESCs cultured in serum-containing medium, the naïve mESCs in 2i medium have reduced basal expression levels of early differentiation genes (Ying et al. 2008; Silva et al. 2008). Unexpectedly, the PRC2-mediated histone H3K27me3, which functions as a major epigenetic mechanism mediating the differentiation gene silencing in mESCs, have been found to have a global reduction in naïve mESCs (Guo et al. 2016; Marks et al. 2012). Previous studies suggested the reduced PRC2 occupancy could be caused by relocation of PRC2 from CGIs to non-CGI DNA demethylated regions in naïve mESCs, or reduced chromatin accessibility to PRC2 at bivalent promoters in the ERK1/ERK2 signaling-deficient mESCs (Walter et al. 2016; van Mierlo et al. 2019; Kumar and Elsasser 2019; Tee et al. 2014).

For the first time, this study provides compelling evidence to demonstrate that the genome-wide decreased PRC2 occupancy at CGIs and bivalent promoters in naïve mESCs are mainly caused by the reduced JARID2-mediated PRC2 recruitment. First, the RNA-seq, qRT-PCR, and WB analyses show that the *Jarid2* expression is significantly reduced in naïve mESCs, while other Polycomb components have similar expression levels between mESC-S and mESC-2i (Fig. 1, S1). JARID2 is a known factor to recruit PRC2 to CGIs in mammalian cells (Peng et al. 2009; Pasini et al. 2010; Shen et al. 2009). Therefore, the decreased *Jarid2* expression provides an important molecular basis for the global reduced PRC2 occupancy in naïve mESCs. Second, the FGF/ERK signaling positively regulates the *Jarid2* expression since the reactivation of FGF/ERK signaling in naïve mESCs upregulates the *Jarid2* expression at both mRNA and protein levels (Fig. 2, S2). Third, similar to the chemical inhibition of ERK signaling in the 2i medium, the genetic disruption of ERK signaling by deletion of *Erk1* and *Erk2* largely reduces the *Jarid2* expression in the mESCs cultured in the serum-containing medium, which can be rescued by the ectopic expression of wild-type *Erk2*, further confirming that ERK signaling positively regulates the expression of *Jarid2* (Fig. 3, S3). Fourth, the ChIP-seq analyses show the global reduced EZH2 occupancy and histone H3K27me3 modification at CGIs and bivalent promoters corelate with the global reduced JARID2 occupancy, which can be fully rescued by the ectopic expression of *Jarid2* in both naïve and *Erk1/Erk2*-dKO mESCs (Fig. 4-6, S4-6). All these results strongly indicate the FGF/ERK signaling determines the global PRC2 occupancy at CGIs and bivalent promoters mainly through the regulation of *Jarid2* expression in mESCs.

Several recent studies have reported that although the H3K27me3 modification is reduced at CGIs, the overall abundance of PRC2 and H3K27me3 in naïve mESCs is increased (Kumar and Elsasser 2019; van Mierlo et al. 2019). Furthermore, the H3K27me3 is redistributed to non-CGI euchromatin and heterochromatin regions, which correlates with the local CpG density (van Mierlo et al. 2019). Since naïve mESCs have global reduced DNA methylation, these results suggest the reduced PRC2 occupancy at CGIs in naïve mESCs could be caused by the relocation of PRC2 from CGIs to new DNA demethylated regions including pericentric heterochromatin (van Mierlo et al. 2019). Although our results do not reveal significant increased EZH2/JARID2 occupancy and H3K27me3 modification at extended 50kb CGI-flanking regions (Fig. S4C-E), our study does not directly conflict with and preclude these conclusions due to the practical difficulty in mapping ChIP-seq reads to heterochromatin. To monitor whether there exists Polycomb redistribution in response to the global DNA demethylation in naïve mESCs, we include KDM2B in our ChIP-seq assay since its CxxC-ZF domain has a high binding affinity to unmethylated CG-rich DNA such as CGIs, and importantly it is known to bind DNA demethylated pericentric heterochromatin in the DNA methyltransferases triple knockout (DNMT-TKO) cells (He et al. 2013; Blackledge et al. 2010; Cooper et al. 2014). However, our study finds both mESC-S and mESC-2i have the same KDM2B occupancy at CGIs (Fig. 4, S4). Since *Kdm2b* has the same expression level in mESC-S and mESC-2i (Fig. S1), the results suggest that KDM2B is unlikely to have a significant redistribution or to mediate PRC2 relocation to non-CGI demethylated regions in naïve mESCs. In addition, the histone H3K4me3 modification at CGIs, what is mainly mediated by MLL2 that binds to CGIs through a similar CxxC-ZF domain as KDM2B (Denissov et al. 2014; Long, Blackledge, and Klose 2013), is increased at CGIs in naïve mESCs (Fig. S4), further arguing against that the CxxC-ZF domain-containing proteins like KDM2B are relocated from CGIs to the new DNA demethylated regions. Finally but importantly, the ectopic expression of *Jarid2* in naïve mESCs fully rescues the global PRC2 occupancy and the histone H3K27me3 modification at CGIs (Fig. 5, S5), further supporting that the reduced PRC2 occupancy at CGIs in naïve mESCs is mainly caused by the reduced JARID2-mediated PRC2 recruitment.

A previous study has reported that ERK1/ERK2 determine the PRC2 occupancy at bivalent promoters through regulating local chromatin accessibility (Tee et al. 2014). Consistent with this report, we find that the global PRC2 occupancy and H3K27me3 modification are reduced in the *Erk1/Erk2*-dKO mESCs, which is rescued by ectopic expression of wild-type *Erk2* (Fig. 6). In addition, we also find that same as naïve mESCs, *Erk1/Erk2*-dKO mESCs have significant reduced *Jarid2* expression, which can be rescued by ectopic expression of wild-type *Erk2* (Fig. 3, S3). Importantly, the ectopic expression of *Jarid2* fully restores the EZH2/JARID2 occupancy and H3K27me3 modification at both CGIs and bivalent promoters in the *Erk1/Erk2*-dKO mESCs (Fig. 6, S6). Since majority of bivalent promoters is associated with CGIs, our study suggests that the reduced JARID2-mediated PRC2 occupancy at CGIs play a more general role leading to the reduction of PRC2 occupancy at bivalent promoters in the *Erk1/Erk2*-depeleted mESCs. However, it is worth pointing out that this study does not preclude the previous conclusion that ERK1/ERK2 also regulate the PRC2 occupancy at bivalent promoters through regulating local chromatin accessibility to PRC2 (Tee et al. 2014).

The Polycomb depletion leads to de-repression of its regulated bivalent genes in mESCs cultured in serum-containing medium, in which mESCs express low-level transcription factors associated with FGF/ERK signaling (Illingworth et al. 2016; He et al. 2013), suggesting Polycomb binding to promoters is essential for setting high transcriptional thresholds to prevent stochastic transcription (Laugesen, Hojfeldt, and Helin 2016). However, previous studies showed that majority of bivalent genes remains silenced although the PRC2 occupancy at their promoters are reduced in naïve mESCs (Marks et al. 2012; Silva et al. 2008; Riising et al. 2014; Galonska et al. 2015). To solve this unexpected discrepancy, our study separates two groups of bivalent genes with distinct transcriptional states, either downregulated or de-repressed, in naïve mESCs (Fig. 7A, S7A, S7B). Since all bivalent promoters have the same reduced PRC2 occupancy in mESC-2i (Fig. S5E-G), the distinct transcriptional status of these two gene groups suggests that the transcriptional activation of bivalent genes is determined by factors other than the PRC2 occupancy at their promoters in naïve mESCs. Further analyses reveal that the downregulated gene group contains multiple primitive endoderm-specific genes known to be activated by the FGF signaling (Chazaud et al. 2006; Hamilton et al. 2019; Illingworth et al. 2016), while the de-repressed gene group includes all known direct targets of Wnt/beta-catenin signaling, suggesting the de-repression of these genes is induced by transcription factors associated with the activated Wnt/beta-catenin signaling in the 2i medium (Fig. 7B-E, S7C, S7D) (Roose et al. 1999; Yan et al. 2001; Lustig et al. 2002; ten Berge et al. 2008; Horvay et al. 2011; Lickert et al. 2000). Finally, more than half de-repressed genes in naïve mESCs, including all known Wnt/beta-catenin direct target genes, are re-silenced after inactivation of Wnt/beta-catenin signaling by removal of GSK3 inhibitor from the 2i medium (Fig. 7C-E), further supporting that the presence of transcription factors, but not the reduced PRC2 occupancy at promoters, plays a predominant role in activating of bivalent genes in naïve mESCs. Of note, this result is consistent with previous reports that the promoter-bound PRC2 does not actively repress transcription but rather sets high transcriptional thresholds for gene activation in cells (Riising et al. 2014; Laugesen, Hojfeldt, and Helin 2016).

In summary, our study provides compelling evidence to reveal a key molecular mechanism by which the FGF/ERK signaling regulates the global PRC2 occupancy at CGIs in naïve mESCs, and to elucidate a fundamental question regarding the function of transcription factors and PRC2-mediated epigenetic mechanisms in transcriptional regulation. Based on the evidence, we propose that the FGF/ERK signaling positively regulates the *Jarid2* expression and promotes the JARID2-mediated PRC2 recruitment to CGIs in mESCs cultured in serum-containing medium. The PRC2 binding to promoters increases the overall transcriptional thresholds, which is essential for preventing the stochastic transcriptional activation of lineage differentiation genes induced by the FGF/ERK signaling in the serum-containing medium. In naïve mESCs, the chemical inhibition of FGF/ERK signaling reduces the *Jarid2* expression and leads to a global reduction of JARID2-mediated PRC2 recruitment to CGIs and bivalent promoters, which consequently reduces the overall transcriptional thresholds of bivalent genes. However, in the absence of FGF/ERK signaling-associated transcription factors, the reduced PRC2 occupancy at promoters alone is insufficient to activate FGF/ERK signaling target genes. In contrast, the Wnt/beta-catenin signaling target genes are de-repressed by the activated Wnt/beta-catenin signaling and its-associated transcription factors in naïve mESCs (Fig. 8)

**Figure 8.**
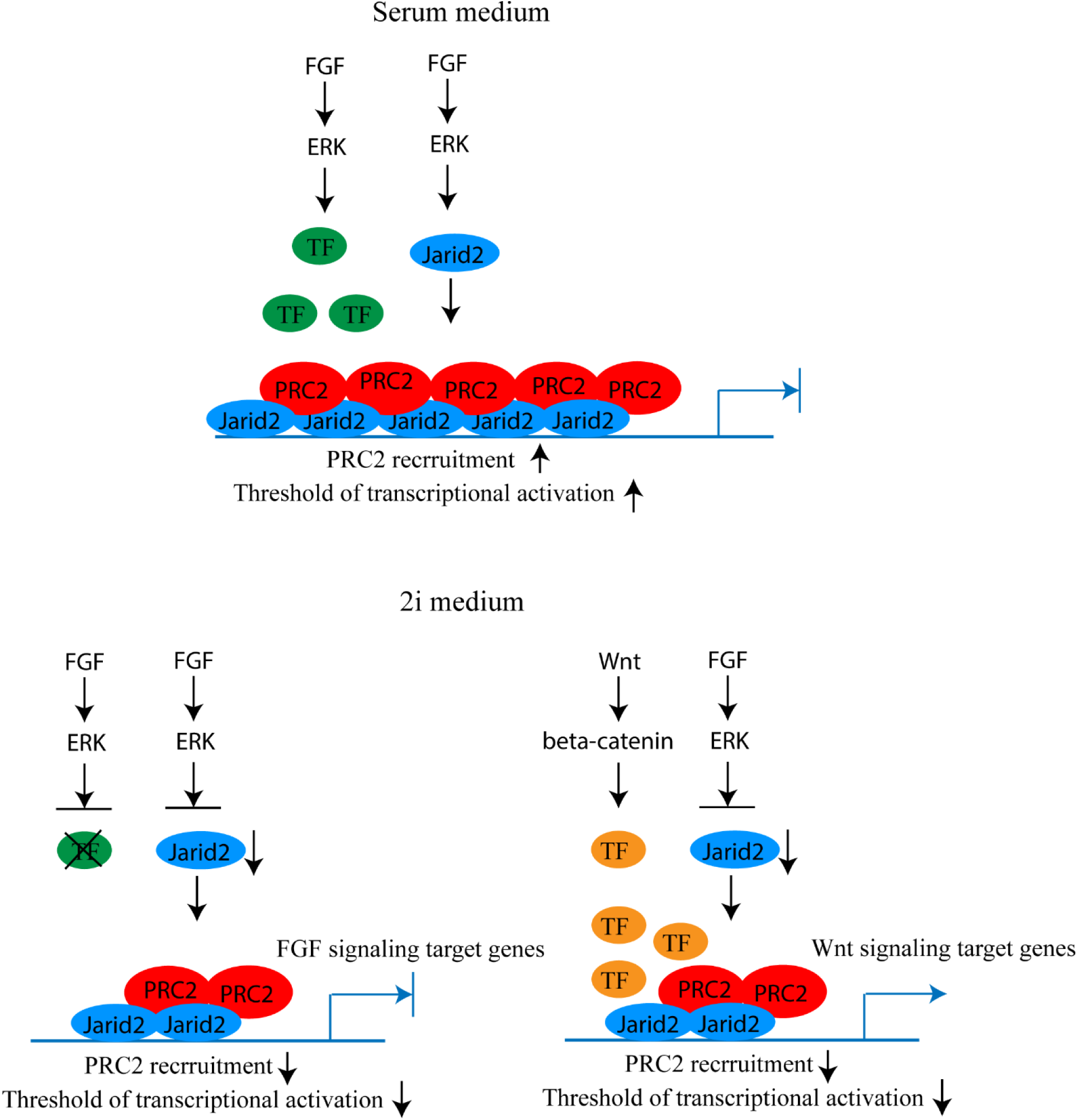
Proposed model of cell signaling coordinating PRC2 recruitment and gene expression in mESCs. In serum-containing medium, FGF/ERK signaling increases the *Jarid2* expression and the JARID2-mediated PRC2 recruitment to CGIs and bivalent promoters, which consequently increases the threshold of transcriptional activation (upper panel). In 2i medium, the deficient FGF/ERK signaling decreases the *Jarid2* expression and JARID2-mediated PRC2 recruitment to CGIs and bivalent promoters, which consequently decreases the threshold of transcriptional activation. In the absence of FGF/ERK signaling-associated transcriptional factors, the FGF/ERK signaling target genes remain silenced. In contrast, the transcription factors associated with the activated Wnt/beta-catenin signaling de-repress the Wnt signaling target genes (lower panel).

## Materials and Methods

### Mouse embryonic stem cell culture

The wild-type E14 mESC line and wild-type E14 mESCs with 3xFlag knock-in to the endogenous Kdm2b gene (He et al. 2013) were maintained on the 0.1% gelatin coated plates in “2i” medium that contained 50% Neurobasal medium (Life Technologies) and 50% DMEM/F12 medium (Life Technologies) supplemented with 1x N2-supplement (Life Technologies), 1x B27 supplement (Life Technologies), 7.5% bovine serum albumin (Life Technologies), 1x GlutaMAX (Life Technologies), 1x beta-mercaptoethanol (Life Technologies), 1mM PD03259010, 3mM CHIR99021, 1000 units/ml leukemia inhibitory factor (ESGRO, EMD Millipore), and 100U/ml penicillin/streptomycin (Life Technologies). The serum-containing mESC culture condition included DMEM medium (Life Technologies) supplemented with 100U/ml penicillin/streptomycin (Life Technologies), 15% fetal bovine serum (Sigma), 1x nonessential amino acid, 1x sodium pyruvate (Life Technologies), 1x GlutaMAX (Life Technologies), 1x beta-mercaptoethanol (Life Technologies) and 1000 units/ml leukemia inhibitory factor (ESGRO, EMD Millipore).

### Western blot analysis

Total proteins were extracted by RIPA buffer and separated by electrophoresis by 8-10% PAGE gel. The protein was transferred to the nitrocellulose membrane and blotted with primary antibodies. The antibodies used for Western Blot and IP-Western Blot analyses included: rabbit anti-FLAG (1:2000, Cell Signaling Technology); rabbit anti-EZH2 (1:1000 Cell Signaling Technology); rabbit anti-RING1B (1:1000, Cell Signaling Technology); rabbit anti-SUZ12 (1:1000 Cell Signaling Technology); rabbit anti-JARID2 (1:1000, Cell Signaling Technology); rabbit anti-p42/44 MAPKs (1:1000 Cell Signaling Technology); rabbit anti-phospho-p42/44 MAPKs ((1:1000 Cell Signaling Technology), rabbit anti-MTF2 (1:1000, Proteintech), rabbit anti-TUBULIN (1:2000, Proteintech), and IRDye 680 donkey anti-rabbit second antibody (1: 10000, Li-Cor). The images were developed by Odyssey Li-Cor Imager (Li-Cor).

### RT-qPCR assays

RNA was extracted and purified from cells using QI shredder (Qiagen) and RNeasy (Qiagen) spin columns. Total RNA (1 µg) was subjected to reverse transcription using Iscript reverse transcription supermix (Bio-Rad). cDNA levels were assayed by real-time PCR using iTaq universal SYBR green supermix (Bio-Rad) and detected by CFX386 Touch Real-Time PCR detection system (Bio-Rad). Primer sequences for qPCR are listed in Supplementary Table 3.

### Lentiviral vector generation and infection

The lentiviral system was obtained from the National Institutes of Health AIDS Research and Reference Reagent Program. To generate mouse *Erk2* and *Jarid2* expression vectors, the complementary DNAs were PCR amplified, fused with P2A and puromycin resistant cassette and cloned into the SpeI/EcoRI sites under the EF1α promoter. To generate lentiviral viruses, the transducing vectors pTY, pHP and pHEF1α–VSVG were co-transfected into HEK293T cells. The supernatant was collected at 24, 36 and 48 hours after transfection, filtered through a 0.45 μm membrane and concentrated using a spin column (EMD Millipore). The mESCs were infected with lentiviral vectors at MOI 1.0. 48 hours after infection, the infected cells are selected by adding puromycin into the medium (1μg/ml).

### Crispr-mediated Erk1/Erk2 gene knock-out in mESCs

The mouse *Erk1* gene gRNA (GGTAGAGGAAGTAGCAGATG) and mouse *Erk2* gene gRNA (GGTTCTTTGACAGTAGGTC and CTTAGGGTTCTTTGACAGT) were cloned into pX330 vector obtained from Addgene. The target vector and pEF1a-pac vector were co-transfected (5:1 ratio) to E14 mESCs using Xfect according to the manufacture’s instruction (TaKaRa, Inc). 48 hours after transfection, Puromycin (1μg/ml) was added to the medium to select transfected cells. The individual clones were manually picked and expanded. The correct knockout clones were selected based on the Sanger sequencing on the targeting sites of genomic DNA, cDNA, and Western blot analysis.

### ChIP-Seq sample preparation

For EZH2, JARID2, RING1B, KDM2B, H3K27me3 and H3K4me3 ChIP, Kdm2b 3xFlag knock-in wild-type E14 cells(He et al. 2013) were fixed with 2mM Ethylene glycol bis[succinimidylsuccinate] (Thermo Scientific) for 1 hour, followed by 10 min in 1% formaldehyde and 5 min 0.125 M glycine to sequence the reaction. Cells were lysed in 1% SDS, 10 mM EDTA, 50 mM Tris-HCl (pH 8.0) and the DNA was fragmented to approximately 200-400 bp by sonification (Branson Sonifier 450). Immunoprecipitation was performed with 3 µg rabbit polyclonal anti-FLAG (Sigma), rabbit anti-EZH2 antibody (1:100, Cell Signaling Technology), rabbit anti-JARID2 antibody (1:100, Cell Signaling Technology), rabbit anti-RING1B antibody (1:100, Cell Signaling Technology), 2ug rabbit anti-H3K27me3 antibody (Diagenode), 2ug rabbit anti-H3K4me4 antibody (Diagenode) overnight at 4°C. Antibody bound DNA-proteins were isolated by protein G plus/protein A agarose beads (EMD Millipore), washed, eluted and reverse cross-linked DNA was extracted by phenol/chloroform and precipitated.

### ChIP DNA preparation for HiSeq4000 sequencing

ChIP DNA library was constructed for HiSeq4000 (Illumina) sequencing using NEBNext UltraII DNA library Prep Kit for Illumina (New England BioLabs, Inc) according to the manufacturer’s instructions. Adapter-ligated DNA was amplified by PCR for 12-14 cycles and followed by size selection using agarose gel electrophoresis. The DNA was purified using QIAquick gel extraction kit (Qiagen) and quantified both with an Agilent Bioanalyzer and Invitrogen Qubit. The DNA was diluted to a working concentration of 20nM prior to sequencing. Sequencing on an Illumina HiSeq4000 instrument was carried out by the Genomics Core Facility at Michigan State University.

### ChIP-Seq data analysis

For the ChIP-Seq data analysis, all sequencing reads were mapped to NCBI build 37 (mm9) of the mouse genome using Bowtie2 (Langmead and Salzberg 2012). Mapped reads were analyzed using the MACS program and bound-regions (peaks) were determined using sequencing reads from input DNA as negative controls (Zhang et al. 2008). When multiple reads mapped to the same genomic position, a maximum of two reads were retained. The statistical cutoff used for peak calling was *P*-value< 10^−8^ and >5-fold enrichment over the control. The mapped sequencing reads were normalized as Counts Per Million Reads (CPM). The normalized reads were binned into 50-bp windows along the genome using the bamCoverage of deepTools program and visualized in the IGV genome browser (Ramirez et al. 2014; Robinson et al. 2011). The datasets of CpG islands and Refseq genes of mm9 mouse reference genome were retrieved from the UCSC table bowser. The heatmap and plot of ChIP-seq reads in the 10kb-windows flanking CGIs or Refseq genes were generated using plotHeatmap and plotProfile in the deepTools program. The subset of bivalent promoters was determined by the 2kb promoter regions upstream of transcriptional start sites that contain both H3K27me3 and H3K4me3 using the bedtools program (Quinlan and Hall 2010).

### RNA-seq sample preparation for HiSeq4000 sequencing

RNA was extracted and purified from cells using QI shredder (Qiagen) and RNeasy (Qiagen) spin columns. Total RNA (1 µg) was used to generate RNA-seq library using NEBNext Ultra Directional RNA library Prep Kit for Illumina (New England BioLabs, Inc) according to the manufacturer’s instructions. Adapter-ligated cDNA was amplified by PCR and followed by size selection using agarose gel electrophoresis. The DNA was purified using Qiaquick gel extraction kit (Qiagen) and quantified both with an Agilent Bioanalyzer and Invitrogen Qubit. The libraries were diluted to a working concentration of 10nM prior to sequencing. Sequencing on an Illumina HiSeq4000 instrument was carried out by the Genomics Core Facility at Michigan State University.

### RNA-Seq data analysis

RNA-Seq data analysis was performed essentially as described previously. All sequencing reads were mapped mm9 of the mouse genome using Tophat2(Kim et al. 2013). The mapped reads were normalized to reads as Reads Per Kilobase of transcript per Million mapped reads (RPKM). The differential gene expression was calculated by Cuffdiff program and the statistic cutoff for identification of differential gene expression is q < 0.05 and 1.5-fold RPKM change between samples. The normalized mapped reads (RPKM) of each RNA-seq experiments were binned into 50bp windows along the genome using the bamCoverage of deepTools program and visualized in the IGV genome browser. The heatmap and plot of gene expression were generated using plotHeatmap and plotProile in the deepTools program. The differential expressed gene lists were input into the DAVID Functional Annotation Bioinformatics Microarray Analysis for KEGG pathway and gene ontology enrichment analyses (https://david.ncifcrf.gov/).

## Acknowledgements

We thank Drs. David Arnosti and Amy Ralston for their critical reading of the manuscript. MSU genomics core facility processed the next-generation sequencing. This work was supported by the National Institute Health NIH grant R01GM127431.

## Author contributions

J.H. conceived the project. M.B.A., Y.G. and J.H. designed and performed the experiments. J.H. and G.I.M. performed the sequencing data analysis. J.H. interpreted the data and wrote the manuscript.

## Supplementary Materials

Fig. S1. The *Jarid2* expression is significantly reduced in naïve mESCs.

Fig. S2. FGF/ERK signaling positively regulates the *Jarid2* expression.

Fig. S3. Knockout of *Erk1/Erk2* reduces the *Jarid2* expression in mESCs cultured in serum-containing medium.

Fig. S4. The genome-wide PRC2 occupancy is largely reduced in naïve mESCs.

Fig. S5. Ectopic expression of *Jarid2* restores the global PRC2 occupancy and histone H3K27me3 modification in naïve mESCs.

Fig. S6. Ectopic expression of *Erk2* or *Jarid2* restores the global PRC2 occupancy and histone H3K27me3 modification at bivalent promoters in *Erk1/Erk2*-dKO mESCs.

Fig. S7. De-repression of bivalent genes is determined by the presence of signaling-associated transcription factors but not the status of PRC2 occupancy in naïve mESCs.

Table S1. Result of KEGG pathway analysis on the genes upregulated in response to the reactivation of FGF/ERK signaling.

Table S2. Result of gene ontology analysis on the bivalent genes that are downregulated in naïve mESCs.

Table S3. Result of gene ontology analysis on the bivalent genes that are upregulated in naïve mESCs.

Table S4. Sequences of all primers used in this study.

**Figure S1.**
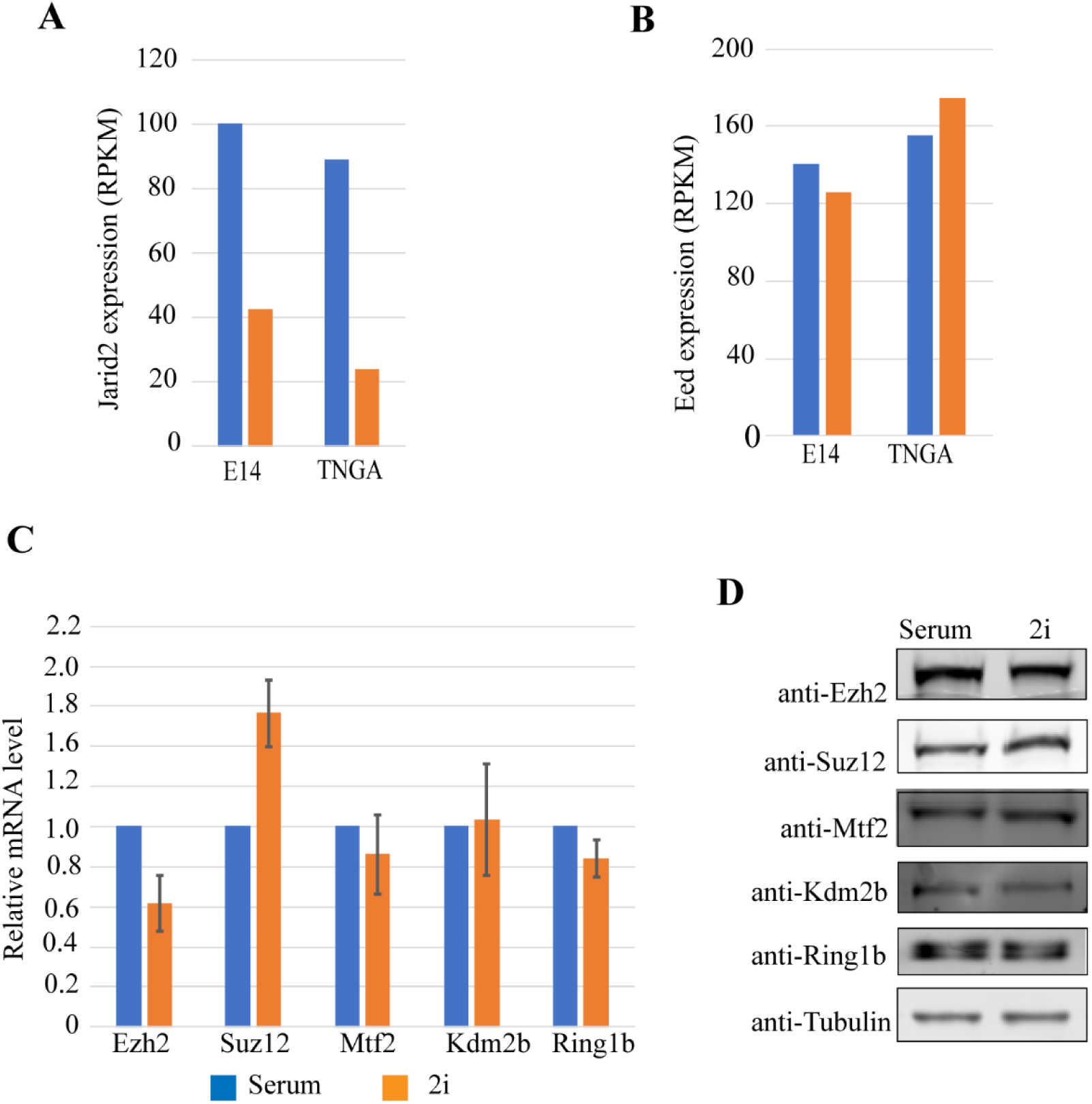
The *Jarid2* expression is significantly reduced in naïve mESCs. **(A)** Re-analysis on a published dataset showing the expression of *Jarid2* in E14 and TNGA mESC lines (Marks et al. 2012). (**B**) Re-analysis on a published dataset showing the expression of *Eed* in E14 and TNGA mESC lines (Marks et al. 2012). (**C**) qRT-PCR analysis showing the expression level of *Ezh2, Suz12, Mtf2, Kdm2b*, and *Ring1b* in mESC-S (serum) and mESC-2i (2i). The results were normalized against levels of Gapdh and the mRNA level in mESC-S was arbitrarily set to 1. The error bars represent standard deviation (n=3). (**D**) Western blot analysis showing the protein level of EZH2, SUZ12, MTF2, KDM2B, and RING1B in mESC-S (serum) and mESC-2i (2i).

**Figure S2.**
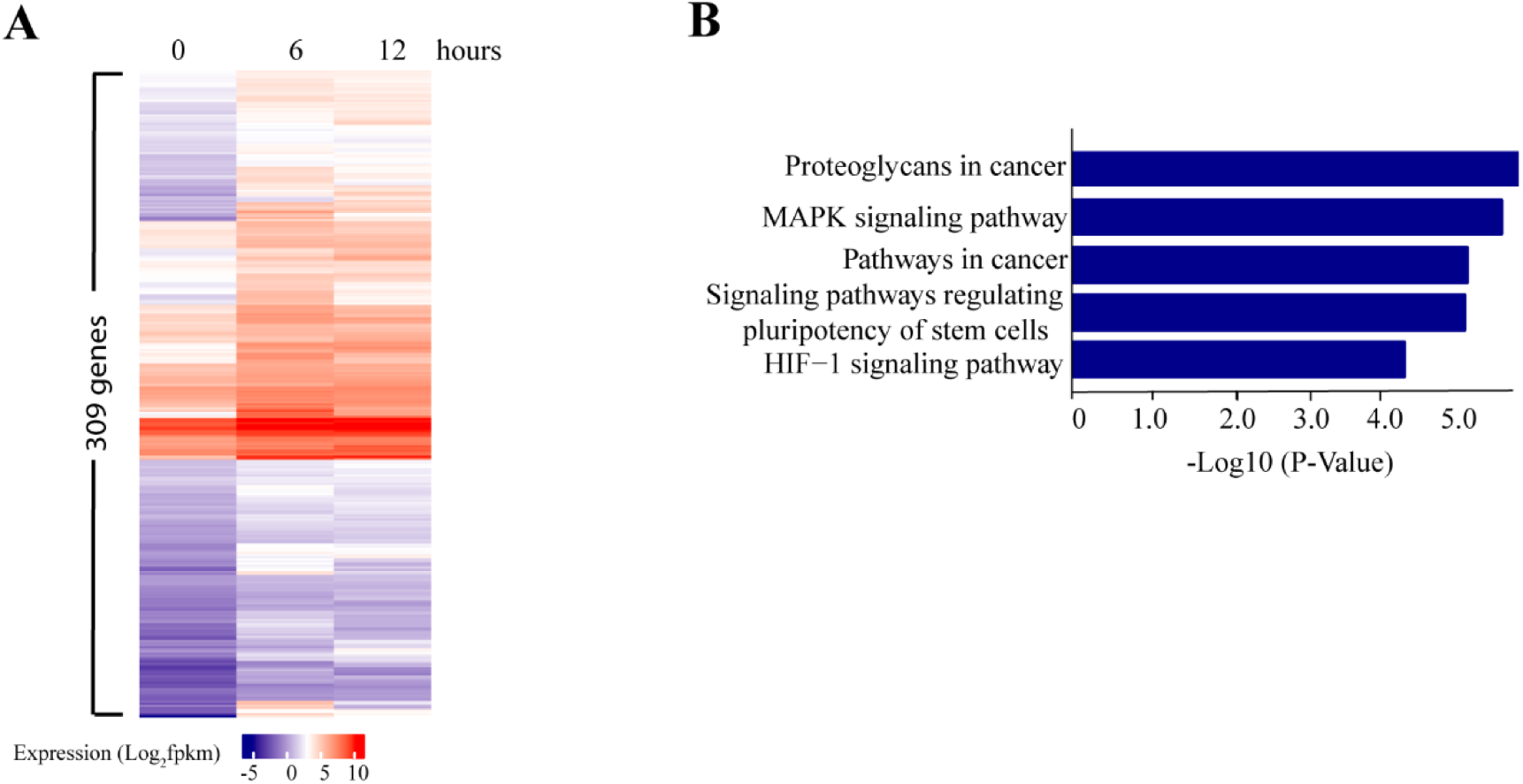
FGF/ERK signaling positively regulates the *Jarid2* expression. (**A**) Heatmap showing 309 genes were upregulated in response to the reactivation of FGF/ERK signaling. (**B**) KEGG signal pathway analysis showing the five top signaling pathways that were activated in response to the reactivation of FGF/ERK signaling (cutoff: Benjamini adjusted p < 0.05).

**Figure S3.**
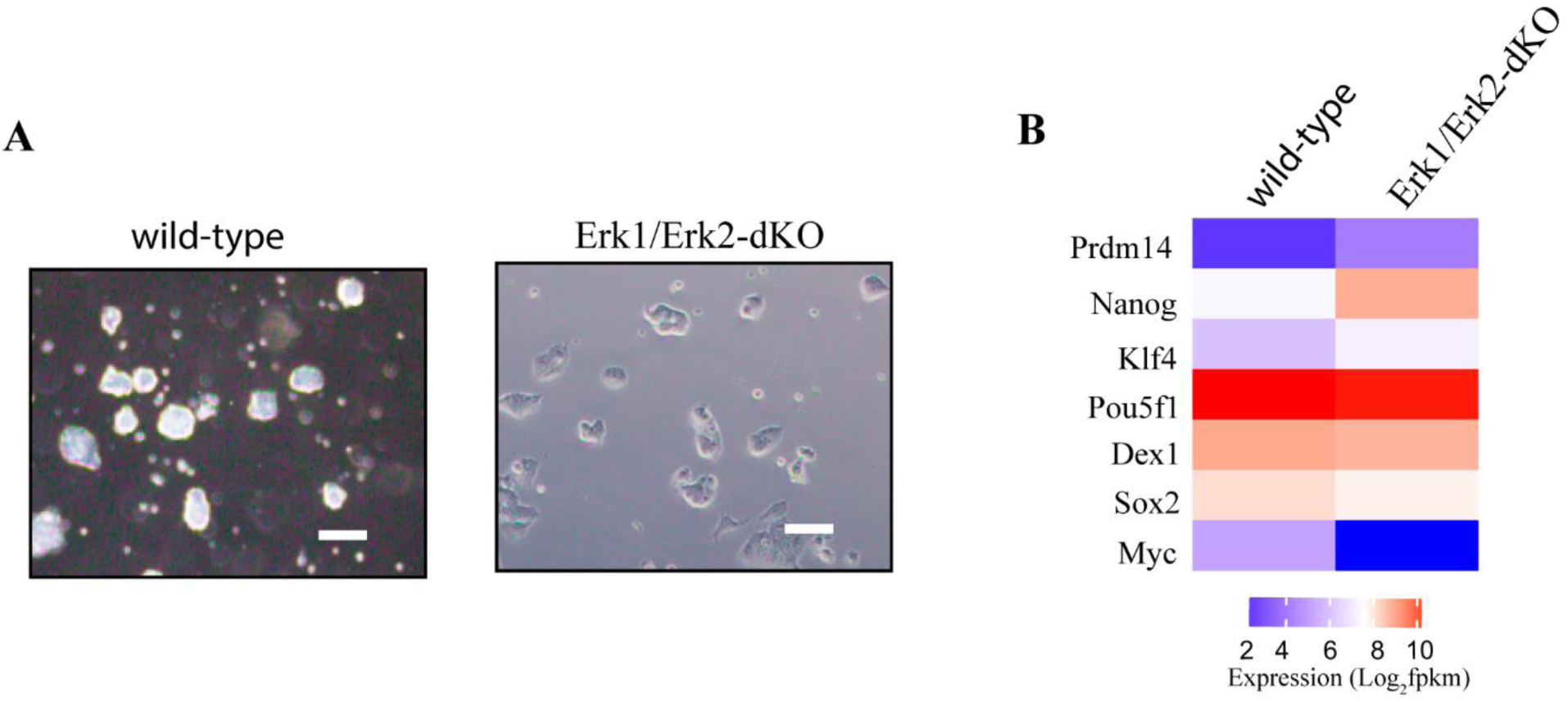
Knockout of *Erk1/Erk2* reduces the *Jarid2* expression in mESCs cultured in serum-containing medium. (**A**) Photos showing the colony morphology of wild-type and *Erk1/Erk2*-dKO mESCs. Scale bar = 100 µm. (**B**) Heatmap showing the expression of pluripotent genes in wild-type and *Erk1/Erk2*-dKO mESCs.

**Figure S4.**
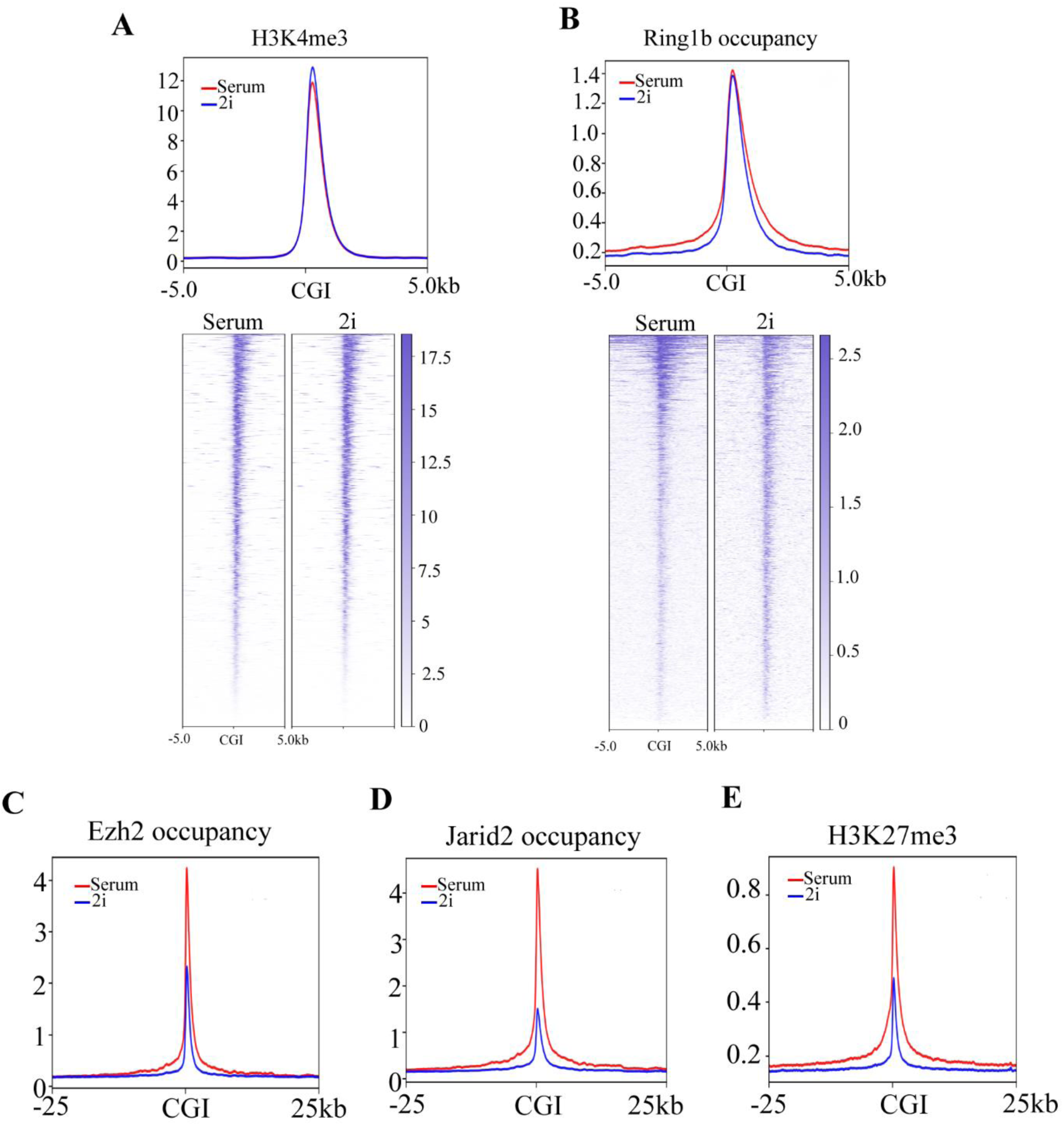
The genome-wide PRC2 occupancy is largely reduced in naïve mESCs. (**A**) The plot (upper) and heatmap (bottom) showing the histone H3K4me3 modification at CGIs and a 10kb-window flanking CGIs in mESC-S and mESC-2i. (**B**) The plot (upper) and heatmap (bottom) showing the RING1B occupancy at CGIs and a 10kb-window flanking CGIs in mESC-S and mESC-2i. (**C**) The plot showing the EZH2 occupancy at CGIs and a 50kb-window flanking CGIs in mESC-S and mESC-2i. (**D**) The plot showing the JARID2 occupancy at CGIs and a 50kb-window flanking CGIs in mESC-S and mESC-2i. (**E**) The plot showing the histone H3K27me3 at CGIs and a 50kb-window flanking CGIs in mESC-S and mESC-2i.

**Figure S5.**
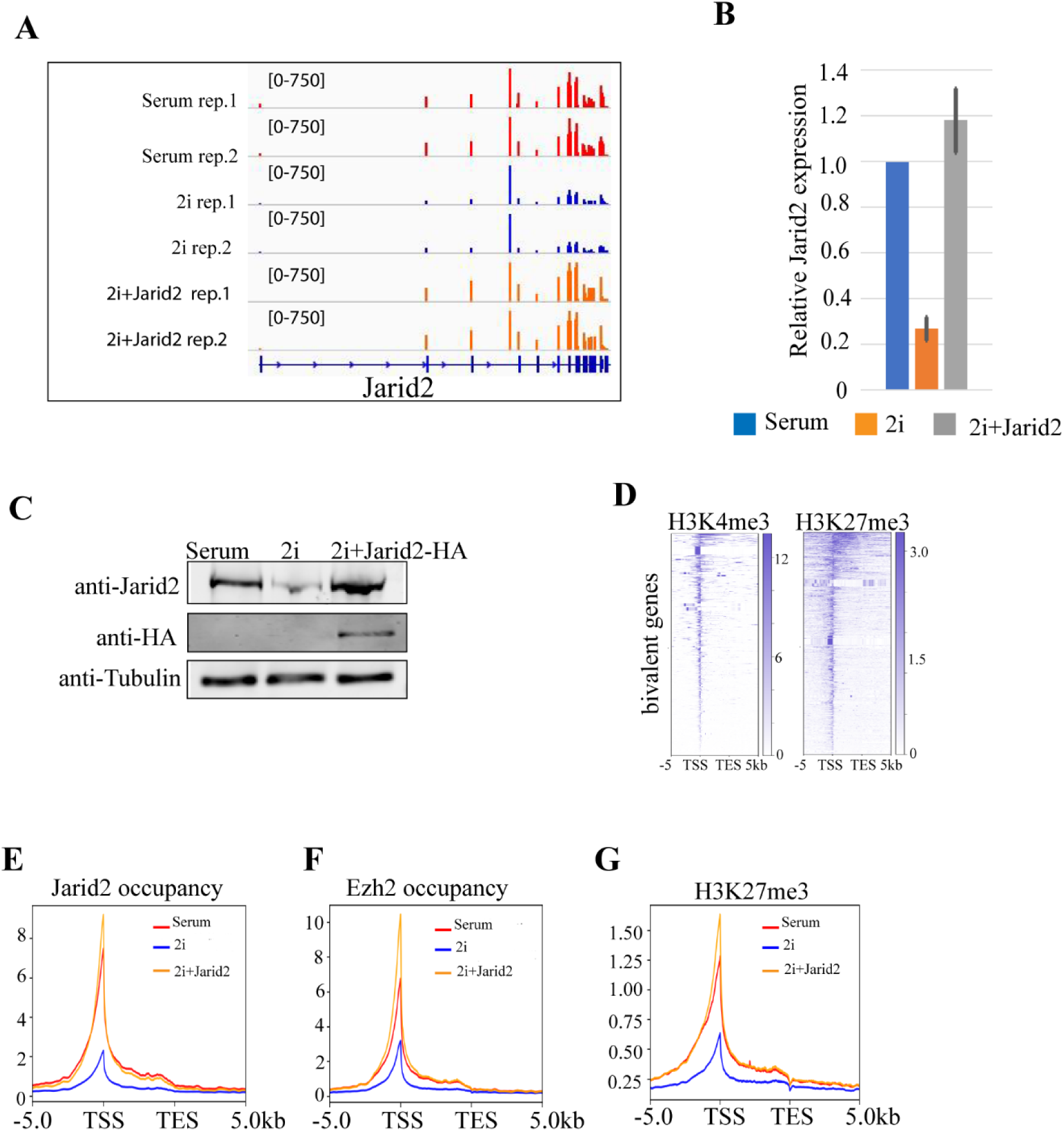
Ectopic expression of *Jarid2* restores the global PRC2 occupancy and histone H3K27me3 modification in naive mESCs. (**A**) The IGV genome browser view of *Jarid2* expression in mESC-S (serum), mESC-2i (2i), and mESC-2i with ectopically expressed *Jarid2* (2i+Jarid2). (**B**) qRT-PCR analysis showing the *Jarid2* mRNA levels in mESC-S (serum), mESC-2i (2i), and mESC-2i with ectopically expressed *Jarid2* (2i+Jarid2). The results were normalized against levels of *Gapdh* and the expression level in mESC-S was arbitrarily set to 1. The error bars represent standard deviation (n=3). (**C**) Western blot analysis showing the JARID2 protein level in mESC-S (serum), mESC-2i (2i), and the mESC-2i with ectopically expressed *Jarid2* (2i+Jarid2). (**D**) The heatmap showing 2830 bivalent gene promoters have both H3K27me3 and H3K4me3 modifications. (**E**) The plot showing the JARID2 occupancy at 2830 bivalent gene promoters in mESC-S (serum), mESC-2i (2i), and mESC-2i with ectopically expressed *Jarid2* (2i+Jarid2). (**F**) The plot showing the EZH2 occupancy at 2830 bivalent gene promoters in mESC-S (serum), mESC-2i (2i), and mESC-2i with ectopically expressed *Jarid2* (2i+Jarid2). (**G**) The plot showing the H3K27me3 modification at 2830 bivalent gene promoters in mESC-S (serum), mESC-2i (2i), and mESC-2i with ectopically expressed *Jarid2* (2i+Jarid2). TSS: transcription starting sites; TES: transcription ending sites.

**Figure S6.**
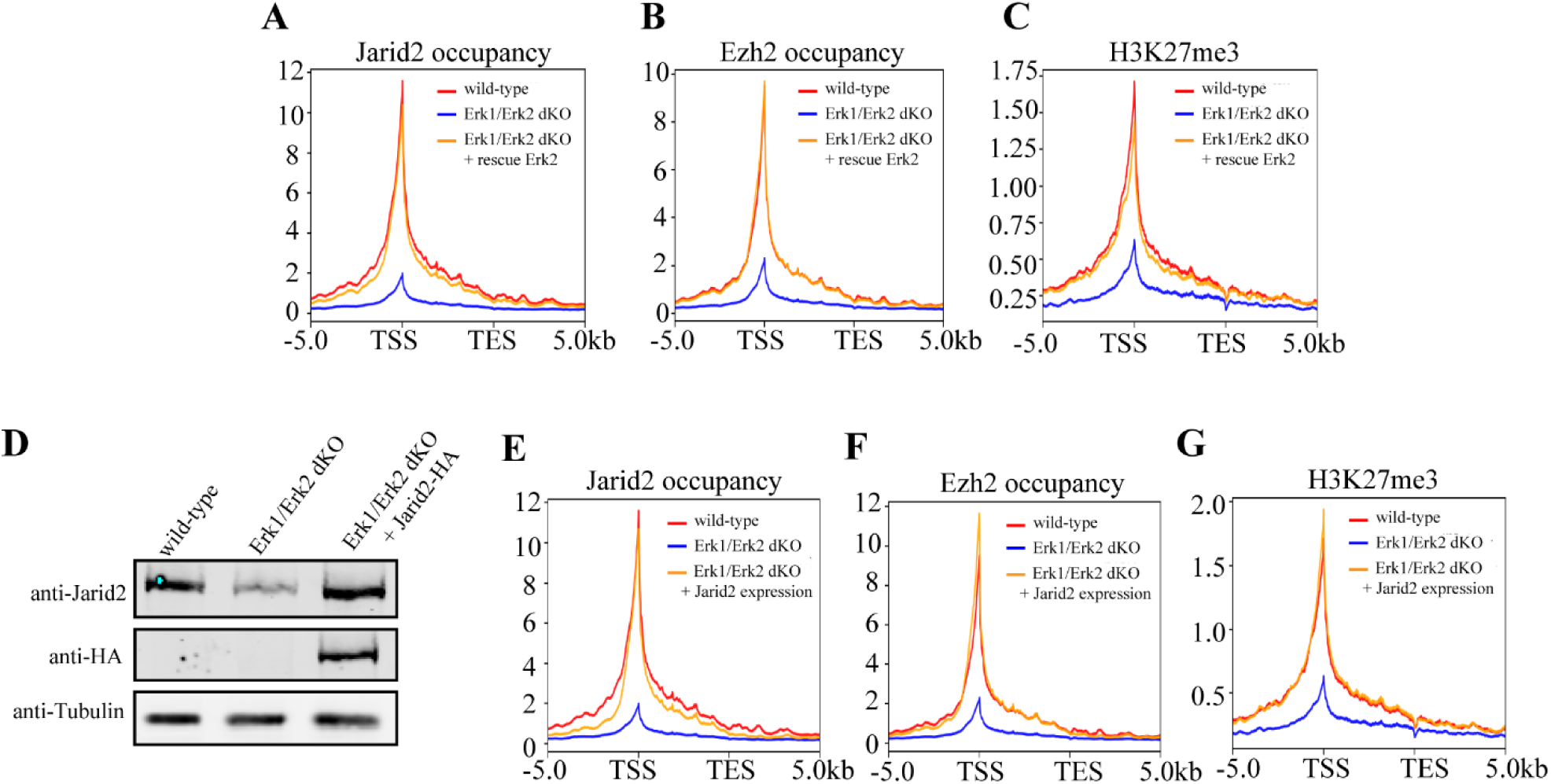
Ectopic expression of *Erk2* or *Jarid2* restores the global PRC2 occupancy and histone H3K27me3 modification at bivalent promoters in *Erk1/Erk2*-dKO mESCs. (**A**) The plot showing the JARID2 occupancy at 2830 bivalent gene promoters in wild-type mESCs (wild-type), *Erk1/Erk2*-dKO mESCs (*Erk1/Erk2*-dKO), and *Erk1/Erk2*-dKO cells rescued with wild-type *Erk2* (Erk1/Erk2-dKO + rescue *Erk2*). (**B**) The plot showing the EZH2 occupancy at 2830 bivalent gene promoters in wild-type, *Erk1/Erk2*-dKO, and *Erk1/Erk2*-dKO + rescue *Erk2* mESCs. (**C**) The plot showing the histone H3k27me3 modification at 2830 bivalent gene promoters in wild-type, *Erk1/Erk2*-dKO, and *Erk1/Erk2*-dKO + rescue *Erk2* mESCs. (**D**) Western blot analysis showing the JARID2 protein level in wild-type mESCs (wild-type), *Erk1/Erk2*-dKO mESCs (*Erk1/Erk2*-dKO), and *Erk1/Erk2*-dKO cells with ectopically expressed *Jarid2* (*Erk1/Erk2*-dKO + *Jarid2* expression). (**E**) The plot showing the JARID2 occupancy at 2830 bivalent gene promoters in wild-type, *Erk1/Erk2*-dKO, and *Erk1/Erk2*-dKO + *Jarid2* mESCs. (**F**) The plot showing the EZH2 occupancy at 2830 bivalent gene promoters in wild-type, *Erk1/Erk2*-dKO, and *Erk1/Erk2*-dKO + *Jarid2* mESCs. (**G**) The plot showing the histone H3k27me3 modification at 2830 bivalent gene promoters in wild-type, *Erk1/Erk2*-dKO, and *Erk1/Erk2*-dKO + *Jarid2* mESCs. TSS: transcription starting sites; TES: transcription ending sites.

**Figure S7.**
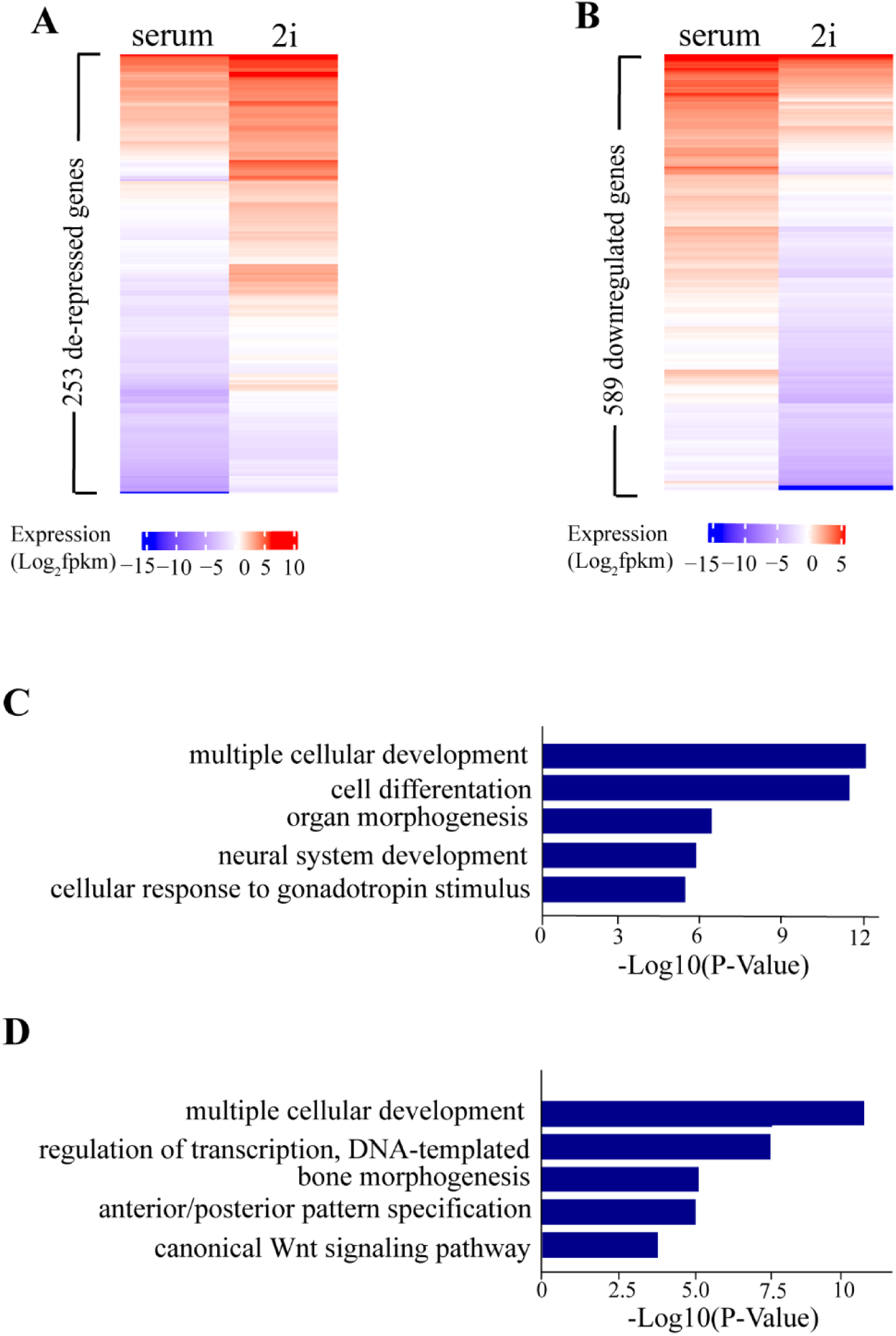
De-repression of bivalent genes is determined by the presence of signaling-associated transcription factors but not the status of PRC2 occupancy in naïve mESCs. (**A**) Heatmap showing 589 bivalent genes downregulated in mESC-2i. (**B**) Heatmap showing 253 bivalent genes de-repressed in mESC-2i. (**C**). Gene ontology analysis showing the enriched GO terms of 589 downregulated bivalent genes in mESC-2i (cutoff: Benjamini adjusted p < 0.05). (**D**) Gene ontology analysis showing the enriched GO terms of 253 de-repressed bivalent genes in mESC-2i (cutoff: Benjamini adjusted p < 0.05).

